# Ultrastructural organization and dynamics of TIRAP filaments

**DOI:** 10.64898/2025.12.13.693976

**Authors:** Arthur Felker, Jan-Hannes Schäfer, Kevin Tanzusch, Marvin Wortmann, Arne Moeller, Jacob Piehler

**Affiliations:** Division of Biophysics, Department of Biology/Chemistry, Osnabrück University, Barbarastraße 11, D-49076 Osnabrück, Germany; Division of Structural Biology, Department of Biology/Chemistry, Osnabrück University, Barbarastraße 11, D-49076 Osnabrück, Germany; Center for Cellular Nanoanalytics, Department of Biology/Chemistry, Osnabrück University, Barbarastraße 11, D-49076 Osnabrück, Germany; Department of Integrative Structural and Computational Biology, Scripps Research; La Jolla, CA, USA

**Author notes:** Correspondence should be addressed to A.F. or J.H.S. These authors contributed equally to this work (Arthur Felker, Jan-Hannes Schäfer).

## Abstract

TIRAP (MAL) is an essential adaptor protein in Toll-like receptor (TLR) signaling, bridging activated receptors to downstream effectors such as MyD88 to initiate pro-inflammatory responses. Assembly of TLR signaling complexes is driven by homotypic interactions between Toll/interleukin-1 receptor (TIR) domains. Although previous studies demonstrated that isolated TIR domains of TIRAP can form filaments *in vitro*, the structural organization and membrane-dependent regulation of full-length TIRAP remained poorly understood. Here, we report a 3.3 Å cryo-electron microscopy (cryo-EM) structure of full-length human TIRAP filaments combined with the first super-resolution imaging of TIRAP assemblies at the plasma membrane of cells. Using complementary lipid-binding assays on supported lipid bilayers (SLBs) and functional live-cell nanopatterning analysis, we identify a critical role of the N-terminal phosphoinositide-binding motif (PBM) in governing filament geometry and assembly dynamics, plasma membrane partitioning, and effector coupling capacity. Collectively, our multimodal analysis provides a comprehensive structure-function framework of TIRAP, highlighting the intricate features of the PBM as a regulator controlling spatiotemporally defined filament assembly and efficient signal induction at the plasma membrane.

## Introduction

Pattern-recognition receptors (PRRs) of the innate immune system detect pathogen-associated molecular patterns (PAMPs) and activate transcriptional responses crucial for host defense and inflammation^1,2^. Among PRRs, Toll-like receptors (TLRs) represent the most prominent family of transmembrane receptors that recognize characteristic pathogen components at the cell surface^3,4^. Ligand-induced TLR dimerization facilitates the proximity of the intracellular Toll/interleukin-1 receptor (TIR) domains, thereby enabling the recruitment of cytosolic TIR-containing adaptor proteins and the assembly of multiprotein signaling complexes through cooperative interactions^5–7^. Downstream signaling cascades include the activation of transcription factors such as NF-κB, AP-1, and IRFs, leading to the expression of pro-inflammatory and antimicrobial genes^8–10^. Dysregulation of TLR signaling is associated with a wide spectrum of diseases, including autoimmune and chronic inflammatory disorders, allergies, cancers, and cardiovascular diseases^11,12^.

TIRAP (also known as MyD88-adaptor-like, MAL) is a critical TIR-domain adaptor protein responsible for recruiting MyD88 through TIR-TIR domain interactions, thereby initiating Myddosome complex formation and downstream signaling^13–18^. Structurally, this adaptor comprises two functional domains. The N-terminal polybasic phosphoinositide-binding motif (PBM) mediates plasma membrane localization through interactions with phosphoinositide lipids (PIs). At the same time, the C-terminal TIR domain is responsible for regulating oligomerization and Myddosome assembly^13,19–23^. Recent structural studies have revealed that isolated TIR domains of multiple TLR adaptors including, MyD88, TRIF, TRAM, and the TIR domain of TLR4 itself, assemble into antiparallel, double-stranded helical filaments in a concentration- and temperature-dependent manner^22,24–27^. Cooperative filament assembly provides a putative mechanistic explanation for the rapid and switch-like amplification of signaling observed in innate immune pathways^28,29^. However, a mechanistic understanding of the determinants governing TIRAP filament assembly in the biological context is lacking. In particular, the role of the PBM in regulating nucleation and assembly of TIRAP filaments at membrane surfaces remains largely unclear.

Here, we have tackled this question by combining high-resolution structural studies, *in vitro* reconstitution at artificial membranes, and super-resolution imaging techniques. Using cryo-electron microscopy (cryo-EM), we resolved the structure of full-length TIRAP filaments and compared the oligomeric arrangement to isolated TIR domains. Reconstitution on supported lipid bilayers (SLBs) revealed a critical role of phosphatidylinositol-4,5-bisphosphate (PIP_2_) in regulating TIRAP oligomer nucleation. We validated the physiological relevance of PBM-mediated membrane binding by visualizing TIRAP filaments at the cell plasma membrane with DNA-PAINT super-resolution microscopy, and by analyzing functional coupling with TLR4 complexes by live-cell nanopatterning assays. Our studies uncover an intricate role of the PBM in steering TIRAP oligomerization in a membrane- and receptor dependent manner.

## Results

### Full-length TIRAP reversibly assembles into compact filaments

Human TIRAP comprises an N-terminal PBM (residues 1-88) featuring three conserved di-lysine clusters, and a C-terminal TIR domain (residues 88-221) responsible for homotypic interactions (**Fig. 1**a, Supplementary Fig. 1a, b). To structurally characterize the self-assembly of TIRAP, we recombinantly expressed full-length human TIRAP and characterized its oligomerization behavior *in vitro* (Supplementary Fig. 2a, b). Negative-stain transmission electron microscopy (TEM) confirmed assembly of TIRAP filaments at physiological temperatures (37°C). This temperature-dependent fibril formation is reversible, with filaments disassembling upon dilution or storage at 4°C (Supplementary Fig. 2c, d), consistent with previous observations^24^. In cryo-EM, TIRAP preferentially assembled into paired strings of about 75 Å in diameter with varying lengths (**Fig. 1**b). Two-dimensional (2D) classification revealed well-aligning repetitive features that translated into clear helical layer lines at 6 Å^−1^ in Fourier space (Supplementary Fig. 3f). Helical reconstruction with a twist of −176° and a rise of 16.6 Å resulted in a global resolution of 3.3 Å (**Fig. 1**c, Supplementary Fig. 3a, b).

**Fig. 1:**
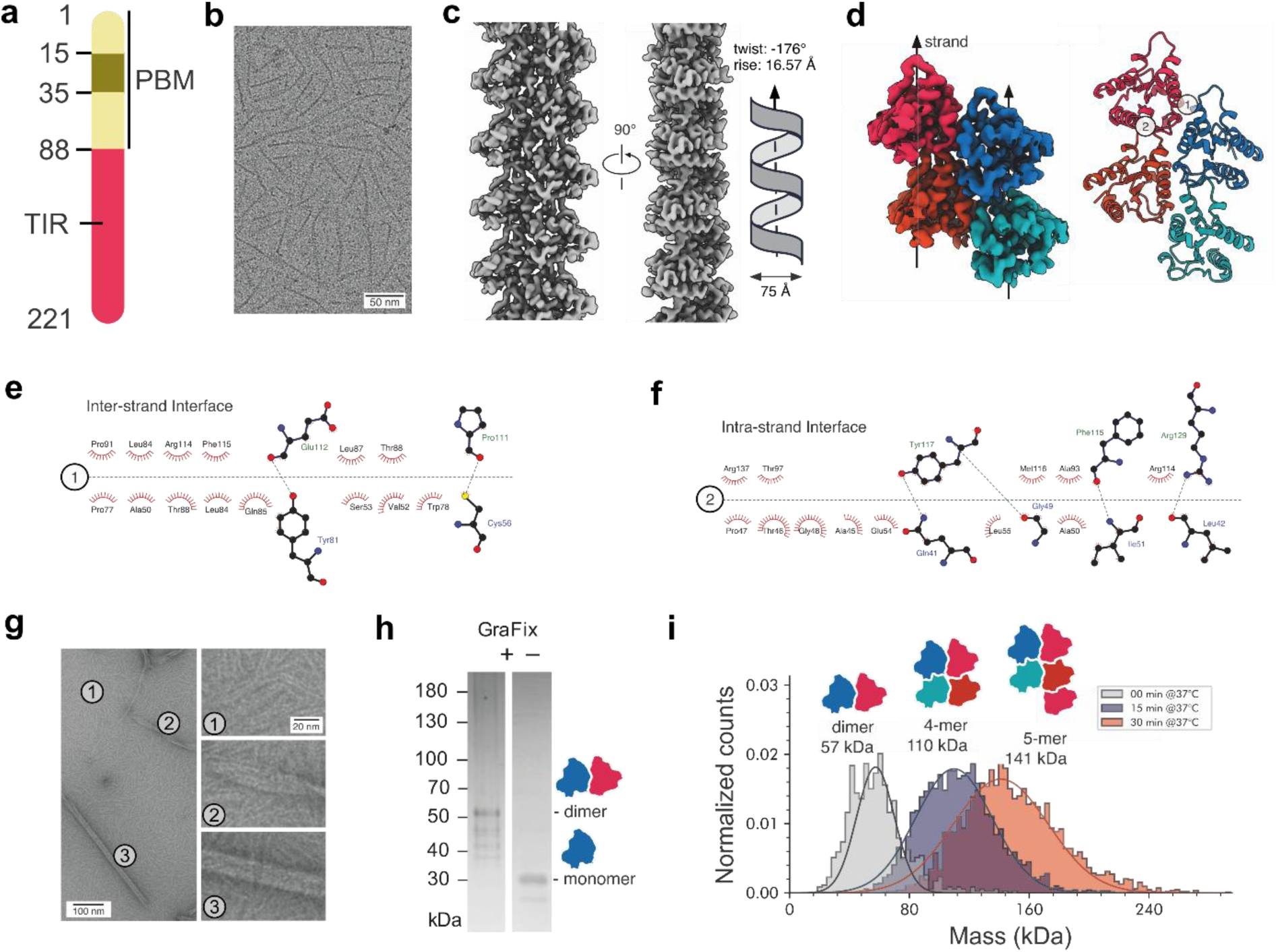
Full-length TIRAP assembles into compact helical filaments via sequential oligomerization. **a** Domain architecture of human TIRAP (UniProt P58753) highlighting the N-terminal PBM (residues 1-88, gradient yellow) containing three conserved di-lysine clusters (positions 15-35) and the C-terminal TIR domain (residues 88-221, red). **b** Representative cryo-EM micrograph of full-length TIRAP filaments. Scale bar: 50 nm. **c** 3.3 Å helical reconstruction at two viewing angles with helical symmetry parameters (twist: −176°, rise: 16.6 Å). **d** Selected density of four TIRAP monomers with strands indicated with arrows and built model (right panel). Numbers indicate contact interfaces (1: inter-strand; 2: intra-strand). **e** Inter-strand interface (interface 1 in d) with key residues labeled. Polar interactions are indicated as green dashed lines and hydrophobic contacts as red half-circles in the 2D interaction diagram (generated with DIMPLOT in LigPlot+)^68^. **f** Intra-strand interface (interface 2 in d). **g** Representative negative-stain EM micrograph showing three assembly morphologies: string-like filaments (1), sheet-like intermediates (2), and tubular structures (3). Insets show magnified views. Scale bars: 100 nm; insets: 20 nm. **h** PAGE fractions of Gradient-Fixation with (+) or without (−) glutaraldehyde cross-linking. **i** Mass photometry analysis of TIRAP oligomerization kinetics at 37°C. Cartoon schematics above peaks indicate oligomeric states. Time points: 0 min (grey), 15 min (blue), 30 min (red). Intensity is given as normalized counts.

Our reconstruction allowed model building of the TIR domain, revealing that TIRAP assembles into pairs of strands through two distinct interfaces (**Fig. 1**d). The inter-strand interface stabilizes the paired architecture through polar interactions between Glu112 and Tyr81, as well as Pro111 and Cys56, complemented by an extensive hydrophobic patch (**Fig. 1**e). The intra-strand interface mediates longitudinal stacking via Tyr117-Gln41, Phe115-Ile51, and Arg129-Leu42 contacts, further stabilized by non-polar interactions (**Fig. 1**f). Notably, the N-terminal PBM was not resolved in our reconstruction. Peripheral fuzzy density observed in 2D class averages (Supplementary Fig. 3f) suggests substantial flexibility of this region in the absence of a membrane environment. This interpretation is consistent with the high predicted disorder of the PBM (Supplementary Fig. 1c) and with NMR data showing that the PBM transitions from a disordered to a helical conformation upon lipid engagement^20^. Importantly, under the conditions used here, full-length TIRAP consistently assembled into compact paired-strand structures, whereas the previously resolved TIR-domain of truncated human TIRAP^TIR^ assembles into large tubular assemblies^24^. This difference suggests that the N-terminal region constrains filament geometry, potentially through steric or electrostatic effects that disfavor higher-order lateral association into tubes.

### Step-wise TIRAP filament assembly into distinct morphologies

To characterize the assembly pathway and intermediates in detail, we analyzed TIRAP by negative-stain TEM, which revealed three distinct morphologies: string-like filaments, sheet-like intermediates, and tubular structures (**Fig. 1**g). The smaller strings dominate the micrographs and may represent an early or stable assembly state. Pairs of strings appear to assemble into sheets and transition into tubes of about 20 nm. The sheet-like and tubular assemblies represent low-abundant species, suggesting they correspond to later stages of higher-order oligomerization. To identify the minimal oligomerization unit, we employed gradient fixation (GraFix)^30^ with glutaraldehyde of assembled TIRAP filaments. PAGE analysis of the cross-linked fractions showed that low-temperature conditions (4°C) only support dimers. In comparison, higher-order oligomers were absent (**Fig. 1**h). We monitored the kinetics of higher-order assembly under physiological conditions (37°C) by mass photometry. TIRAP oligomerization proceeded in a stepwise manner, with dimers detectable immediately, followed by tetramers after 15 minutes and pentamers after 30 minutes (**Fig. 1**i). Prolonged incubation resulted in broader particle size distribution, consistent with the varying filament lengths and morphologies observed by TEM (**Fig. 1**g). Together, these data demonstrate that TIRAP undergoes sequential, temperature-dependent oligomerization initiated from constitutive dimers, ultimately forming extended helical filaments.

### The N-terminal domain of TIRAP regulates lipid binding and filament geometry

Next, we investigated the functional role of the N-terminal PBM in lipid binding and its influence on filament geometry. The PBM contains three conserved di-lysine clusters (Supplementary Fig. 1b) that are predicted to mediate electrostatic interactions with negatively charged phosphoinositides (PIs)^19,20,23^. To characterize the lipid-binding specificity of TIRAP, we performed a lipid overlay assay and subsequent detection of bound His-tagged TIRAP (**Fig. 2**a). We observed TIRAP binding to phosphatidylinositol (PI), phosphatidylinositol 4-phosphate (PI4P), and phosphatidylinositol-4,5-bisphosphate (PIP_2_), indicating recognition of both unphosphorylated and phosphorylated PIs (**Fig. 2**b, left). Additionally, we observed binding to phosphatidylserine (PS) and cholesteryl hemisuccinate (CHS), consistent with a charge-driven interaction. Upon truncating the amino-terminal domain (Δ79), lipid-binding was largely abolished, in line with the hypothesis that the PBM is responsible for membrane recruitment of TIRAP. Interestingly, a residual affinity towards PI4P remained, indicating binding to mono-phosphorylated PIs outside the truncated residue range (**Fig. 2**b, middle). A less severe phenotype was observed by the T28D variant, which only lost sensitivity towards unphosphorylated PI (**Fig. 2**b, right).

**Fig. 2:**
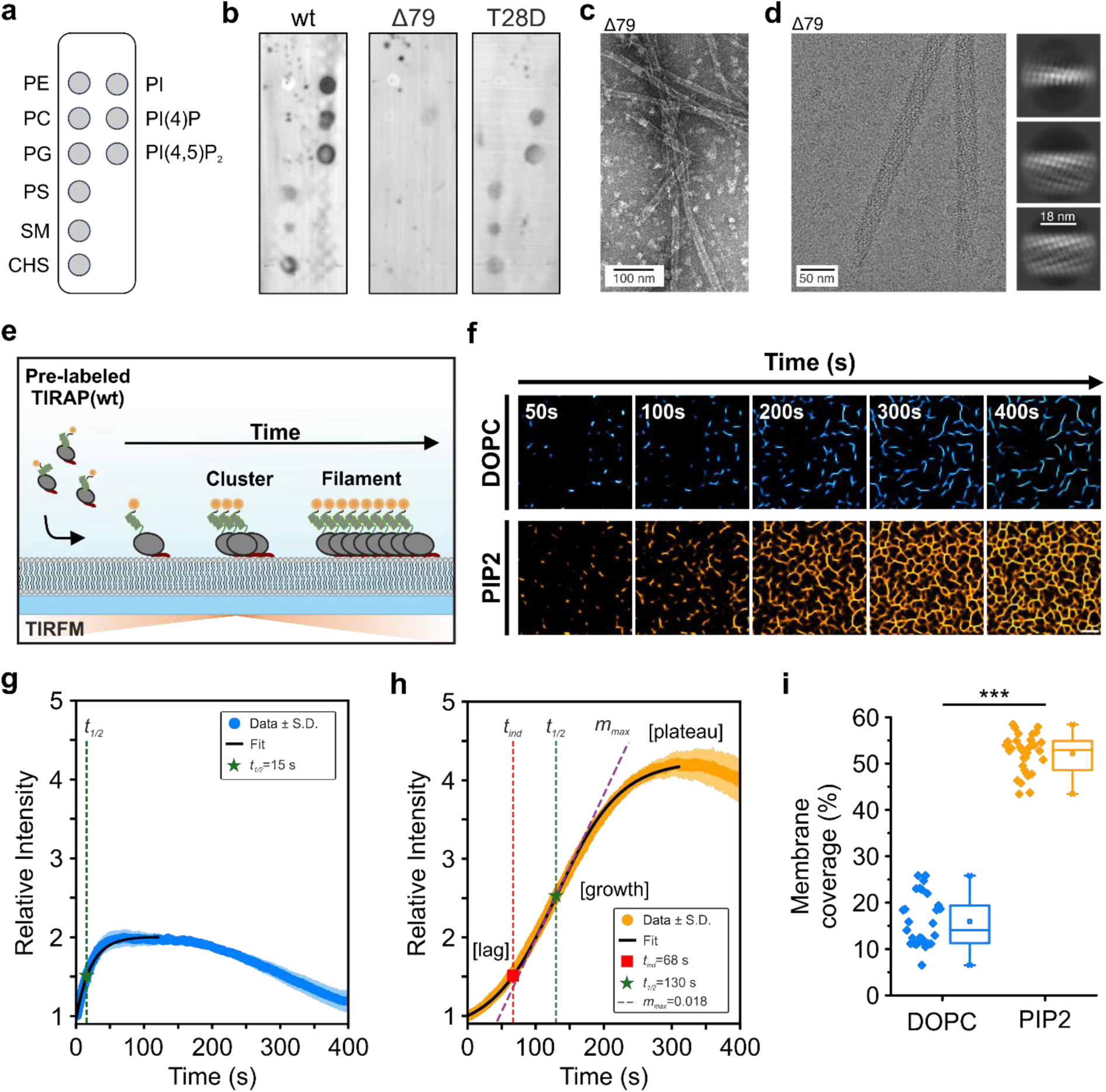
The N-terminal PBM regulates lipid binding and membrane-dependent TIRAP filament assembly. **a** Lipid-overlay assay for lipid binding of His-tagged TIRAP variants. **b** Lipid-binding profiles of wild-type TIRAP (wt), N-terminal deletion mutant (Δ79), and phosphomimetic variant (T28D). **c** Negative-stain EM micrograph of the Δ79 TIRAP variant. Scale bar: 100 nm. **d** Representative cryo-EM micrograph of Δ79 filaments (left) and selected 2D class averages (right). Scale bars: 50 nm (micrograph); 18 nm (2D classes). **e** Schematic of a time-lapse TIRF microscopy experiment on SLBs. Pre-labeled His_6_-TIRAP (Dy547-trisNTA) is applied to DOPC (100 mol%) or DOPC:PIP_2_ (95:5 mol%) bilayers. **f** Representative time-lapse TIRF images (10 µM TIRAP) on DOPC (blue, top row) and PIP_2_-containing bilayers (orange, bottom row). Scale bar: 2 µm. **g** Kinetics of TIRAP assembly on DOPC membranes. Key parameters are indicated. Blue: Data ± s.d.; black line: exponential fit. **h** Kinetics of TIRAP assembly on PIP_2_ bilayers showing lag phase, growth phase, and plateau. Key parameters are indicated. Orange: Data ± s.d.; black line: Finke-Watzky^60^ fit. **i** Membrane coverage after t = 400 s. Box plot indicates the data distribution of the second and third quartiles (box), median (line), mean (square), and 1.5x interquartile range (whiskers). Each dot represents individual regions of interest (n > 25 ROIs per condition, 2 independent experiments). ***p < 0.001, two-sample Kolmogorov-Smirnov test.

Importantly, the Δ79 variant retained the capacity to form higher-order assemblies. However, negative-stain and cryo-EM revealed that Δ79 TIRAP predominantly formed tubular structures with a diameter of 20 nm (**Fig. 2**c, d). Selected 2D class averages from cryo-EM confirmed the tubular morphology (**Fig. 2**d), although helical reconstruction of these assemblies did not yield interpretable densities, likely due to internal flexibility. Notably, the string-like intermediates that dominated full-length TIRAP assemblies were rarely observed in the Δ79 variant, corroborating the notion that the N-terminal region stabilizes the compact architecture and may regulate the transition between different oligomeric states. These results suggest a functional coupling between the TIR-domain and the PBM, with the latter constraining the geometry to compact paired-strand ribbons rather than extended tubes. Such functional coupling implicates that membrane engagement via the PBM may further regulate TIRAP assembly dynamics in a spatiotemporally controlled manner.

### TIRAP filament assembly on supported-lipid membranes is regulated by PIP_2_

Considering the PBM’s specific lipid affinity, we hypothesized that binding to negatively charged membranes locally concentrates TIRAP and thereby promotes filament formation. To test this, we monitored TIRAP filament assembly on SLBs, composed of either DOPC alone or DOPC with 5 mol% PIP_2_, by time-lapse total internal reflection fluorescence (TIRF) microscopy. Bilayer integrity and lateral fluidity were confirmed by fluorescence recovery after photobleaching (FRAP), using DHPE-OG488 as a fluorescent lipid tracer (Supplementary Fig. 4a). For quantitative analysis of membrane-dependent binding and self-assembly, recombinant His_6_-TIRAP was fluorescently labeled with Dy547-trisNTA loaded with Ni^2+^ ions^31,32^, and applied to the SLBs (**Fig. 2**e). As a control, Dy547-trisNTA was added in the absence of His-tagged protein. Under these conditions, no membrane interaction was observed, and the fluorescence signal remained at background level (Supplementary Fig. 4f), excluding nonspecific binding of the fluorophore-chelator complex to the SLB.

At low TIRAP concentration (1 µM), both membrane compositions supported rapid formation of diffraction-limited membrane-associated puncta (Supplementary Fig. 4b). However, the kinetics of membrane recruitment showed marked differences. On DOPC membranes, TIRAP binding followed simple exponential kinetics (*t*_1/2_ ≈ 17 s), reaching saturation within 120 s (Supplementary Fig. 4c). In contrast, on PIP_2_-containing bilayers TIRAP binding exhibited sigmoidal kinetics with reduced initial slope and a short lag phase (*t_ind_* ≈ 4.7 s), followed by accelerated growth (*t*_1/2_ ≈ 45 s) and sustained protein binding until ∼300 s, reaching a slightly higher final intensity (Supplementary Fig. 4d). Despite enhanced membrane recruitment (Supplementary Fig. 4e), the delayed cluster formation suggests that PIP_2_ engagement imposes kinetic constraints on initial oligomerization.

This effect became more pronounced at 10 µM TIRAP, where both membrane compositions supported a transition from puncta to extended filamentous structures (**Fig. 2**f, Supplementary Fig. 4g). On DOPC membranes, filaments emerged from initial clusters through exponential growth (*t*_1/2_ ≈ 15 s), reaching a plateau intensity of 2.0 (1.7-fold increase over 1 µM conditions) after ∼60 s (**Fig. 2**g). Filaments remained largely isolated with minimal interconnectivity, yet continued to elongate until the end of the time course, likely reflecting ongoing assembly masked by moderate photobleaching. Strikingly, PIP_2_-containing SLBs showed a fundamentally different assembly pathway (**Fig. 2**h). A substantially prolonged lag phase (*t_ind_* ≈ 68 s), representing a ∼14-fold increase compared to 1 µM conditions, preceded rapid filament growth (*t_1/2_* ≈ 130 s) with a maximum assembly slope (*m_max_* = 1.84 × 10^−2^) exceeding that at 1 µM by ∼7.5-fold. Filaments continued to elongate and interconnect until ∼300 s, reaching a significantly higher plateau intensity of 4.2 together with higher membrane coverage (**Fig. 2**f, i). The sharp transition from a prolonged lag phase to rapid cooperative growth aligns with switch-like behavior predicted by signaling-by-cooperative-assembly-formation (SCAF) models^27,28,33^. Together, these results suggest that membrane binding via the PBM enriches TIRAP at the membrane in a oligomerization-incompetent conformation that probably requires binding of monomers from solution to trigger cooperative filament assembly. Detailed kinetic parameters for all conditions are provided in Supplementary Table 2.

### Assembly of TIRAP filaments at the plasma membrane depends on its PBM

Next, we examined the process of filament formation in the cellular environment. To this end, HeLa cells were transiently transfected with mEGFP-tagged wild-type TIRAP (TIRAP-wt; WT) or a deletion mutant lacking the PBM (TIRAP-ΔPBM; ΔPBM), which abolishes phosphoinositide binding (**Fig. 2**b). First, we assessed the impact of TIRAP expression levels on filament assembly by comparing cells based on their mean fluorescence intensity observed by TIRF microscopy when focusing on the plasma membrane (“PM intensity”), covering ∼5-fold differences (**Fig. 3**a, b). At low PM intensity, both TIRAP-wt and TIRAP-ΔPBM showed diffuse localization, with minimal evidence of filamentous structures (**Fig. 3**c, d, Supplementary Fig. 5a, b). In contrast, filaments formed for both variants for cells with high PM intensity (**Fig. 3**c, d, Supplementary Fig. 5a, b). To uncover the structural assembly of TIRAP at the plasma membrane, we turned to DNA-PAINT super-resolution imaging^34,35^ by targeting the mEGFP-tagged TIRAP with a GFP-specific nanobody conjugated to a DNA docking strand (**Fig. 3**a). DNA-PAINT reconstructions confirmed minimal occurrence of filamentous structures at low PM intensity for both variants. However, distinct filament morphologies were observed at high PM intensity (**Fig. 3**c, d, middle and bottom rows, Supplementary Fig. 6a, b), matching our TEM observations *in vitro* (**Fig. 1**g, **Fig. 2**d). Quantitative evaluation of the super-resolved filaments revealed significant differences in size and density that directly reflects the PBM’s structural role. TIRAP-wt assembled into densely arranged networks characterized by a median filament length of approximately 1.2 µm and a median width of about 60 nm (**Fig. 3**e). In contrast, TIRAP-ΔPBM formed significantly longer filaments at approximately 1.7 µm with markedly increased width at about 105 nm, yet their filament density was approximately seven-fold lower compared to TIRAP-wt (**Fig. 3**e, f, Supplementary Fig. 6c), consistent with our TEM data of TIRAP-Δ79 (**Fig. 2**c, d). Averaging aligned filament fragments confirmed these structural differences, revealing a more compact architecture for TIRAP-wt compared to a broader assembly for TIRAP-ΔPBM (**Fig. 3**g), in line with the different geometries identified by cryo-EM. Together, these results highlight an intricate role of the PBM in regulating TIRAP membrane binding and filament assembly in cells in a similar manner as observed *in vitro*.

**Fig. 3:**
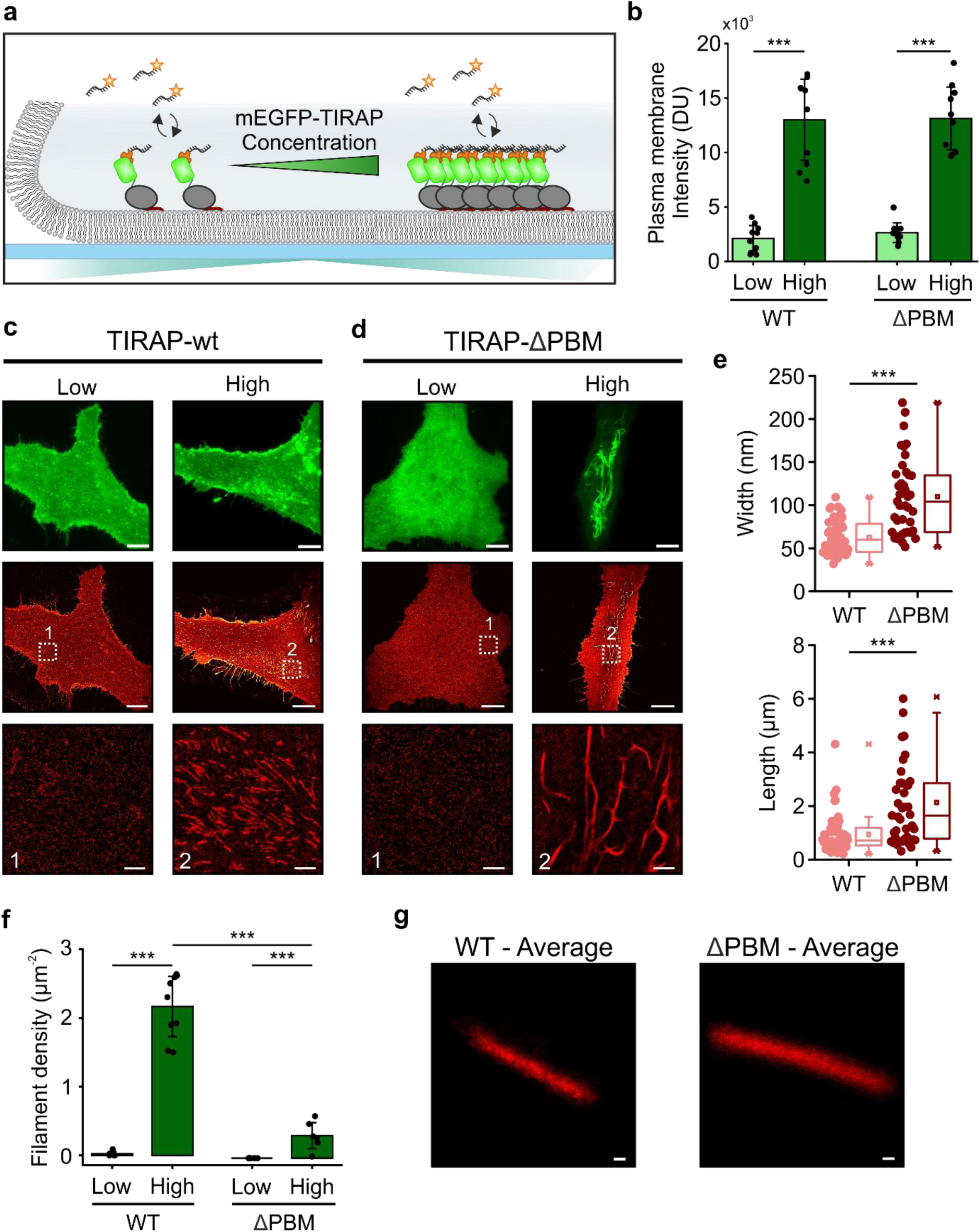
Concentration-dependent TIRAP filament assembly in cells. **a** Schematic illustrating the concentration-dependent lateral self-assembly of mEGFP-TIRAP into filaments. DNA-PAINT imaging was performed by utilizing an anti-GFP nanobody (orange) conjugated with a docking strand for targeted imaging. **b** Plasma membrane (PM) TIRF fluorescence intensities for TIRAP-wt and TIRAP-ΔPBM at low and high expression levels (n = 10 cells per condition; ***p < 0.001, two-sample Kolmogorov-Smirnov test). **c, d** Representative TIRF (top, green) and DNA-PAINT (middle and bottom, red) images for TIRAP-wt (**c**) and TIRAP-ΔPBM (**d**) at low and high PM intensity. Dashed boxes (1, 2) indicate regions enlarged in the bottom row. Scale bars: overview images, 10 µm; DNA-PAINT zoom-in, 1 µm. **e** Filament width (top) and length (bottom) from DNA-PAINT reconstructions at high PM intensity (TIRAP-wt: n = 60 filaments; TIRAP-ΔPBM: n = 40 filaments; ***p < 0.001, two-sample Kolmogorov-Smirnov test). Box plot indicates the data distribution of the second and third quartiles (box), median (line), mean (square), and 1.5x interquartile range (whiskers). **f** Filament density at low and high PM intensity (n > 5 regions of interest per condition; ***p < 0.001, two-sample Kolmogorov-Smirnov test). **g** Averaged DNA-PAINT reconstructions for TIRAP-wt (left) and TIRAP-ΔPBM (right) from 76 and 60 aligned filament segments, respectively. Scale bars: 50 nm.

### The PBM of TIRAP mediates stable plasma membrane recruitment in cells

Having demonstrated that elevated intracellular concentrations of TIRAP promote filament formation in cells (**Fig. 3**), we next investigated their association with the PM in more detail. Confocal laser-scanning microscopy (cLSM) revealed ∼2.4-fold higher PM enrichment for TIRAP-wt compared to TIRAP-ΔPBM, which accumulated predominantly in the cytosol (Supplementary Fig. 7a, b), consistent with impaired PM binding capacity of the deletion mutant. To unambiguously assess stable PM interaction while excluding cytosolic contributions, we generated polymer-supported plasma membranes (PSPMs)^36^. In this approach, intact PM from HeLa cells co-expressing mEGFP-TIRAP and a transmembrane anchor (HaloTag-mTagBFP-TMD) is covalently tethered to PLL-PEG-HTL-coated surfaces via the extracellular HaloTag protein (**Fig. 4**a). Following membrane anchoring, cells were treated with Latrunculin B to disrupt the actin cytoskeleton, allowing subsequent mechanical removal of cell bodies while leaving the isolated PM stably immobilized on the surface. Successful PSPM generation was validated by differential interference contrast (DIC) microscopy, confirming the absence of cell bodies, and by homogeneous distribution of the HaloTag-mTagBFP-TMD tethering construct (**Fig. 4**b, Supplementary Fig. 8). TIRF microscopy revealed an approximately 30-fold higher membrane-associated fluorescence for TIRAP-wt compared to TIRAP-ΔPBM (**Fig. 4**e), demonstrating the essential role of the PBM for stable PM association in cells.

**Fig. 4:**
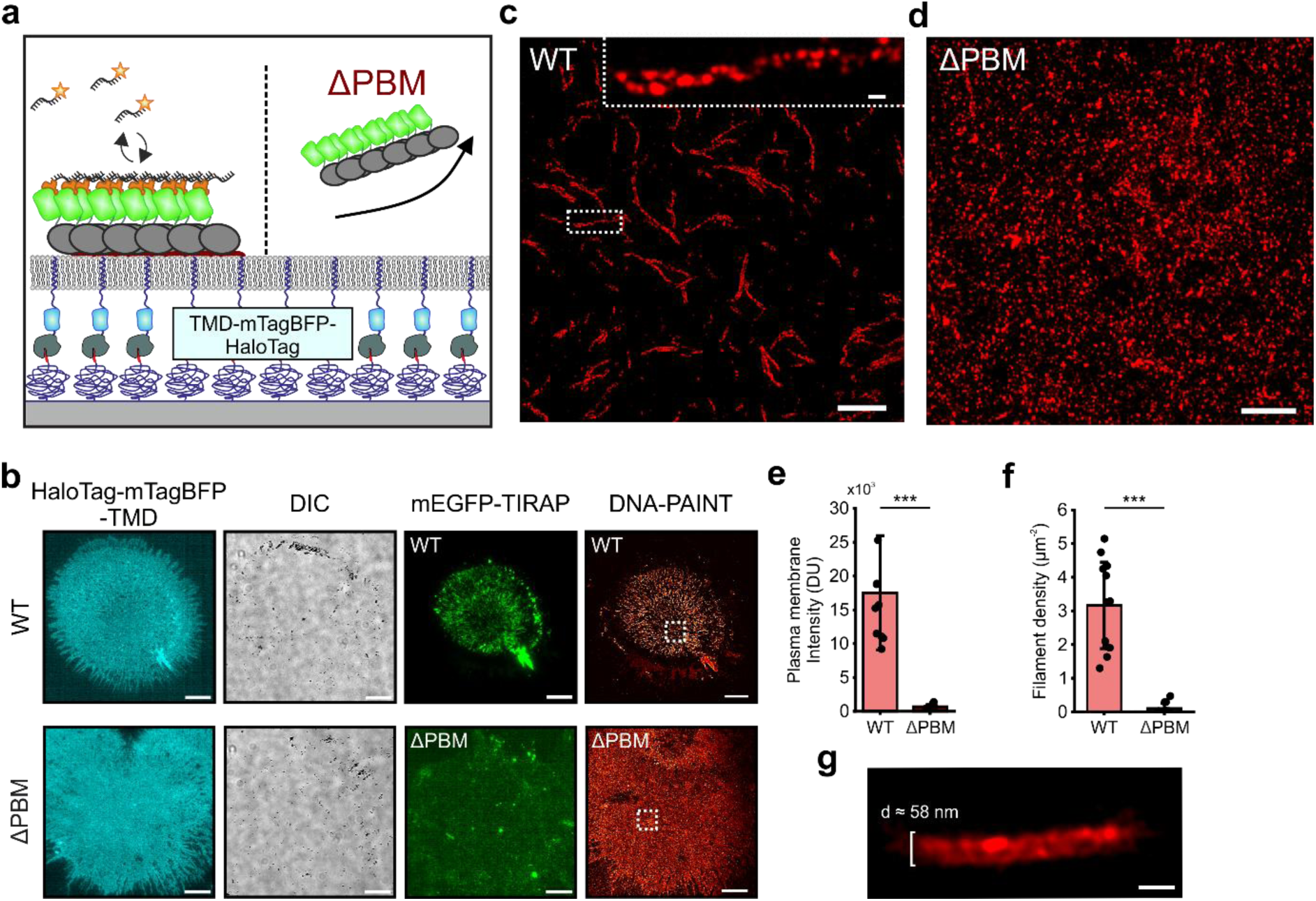
Plasma membrane binding of TIRAP filaments depends on the PBM. **a** Schematic of the proposed differences in PM interaction of TIRAP-wt (left) and TIRAP-ΔPBM (right) on polymer-supported plasma membranes (PSPMs). PMs from HeLa cells expressing mEGFP-TIRAP (green) and HaloTag-mTagBFP-TMD anchor (cyan) are covalently tethered to PLL-PEG-HTL-coated surfaces. Membrane-associated TIRAP filaments were imaged by DNA-PAINT utilizing an anti-GFP nanobody (orange) conjugated with a docking strand. **b** Representative TIRF images of PSPMs from TIRAP-wt (top row) and TIRAP-ΔPBM (bottom row) expressing cells. Left: HaloTag-mTagBFP-TMD anchor (cyan). Middle left: DIC confirming absence of cell bodies. Middle right: mEGFP-TIRAP (green). Right: DNA-PAINT super-resolution reconstruction (red). Scale bars: 10 µm. **c, d** Magnified DNA-PAINT reconstructions from regions indicated by white dashed boxes in **b** for TIRAP-wt (**c**) and TIRAP-ΔPBM (**d**). Upper inset in **c** show individual filament segment. Scale bars: 1 µM; zoom-in, 50 nm. **e, f** Quantification of membrane-associated mEGFP-TIRAP fluorescence intensity (**e**) and filament density from DNA-PAINT reconstructions (**f**) on PSPMs (TIRAP-wt: n = 10 PSPMs; TIRAP-ΔPBM: n = 10 PSPMs; ***p < 0.001, two-sample Kolmogorov-Smirnov test). **g** Averaged DNA-PAINT reconstruction of TIRAP-wt filaments on PSPMs from 67 aligned segments. Scale bar: 50 nm.

To visualize filament organization on PSPMs, we again employed DNA-PAINT super-resolution microscopy (**Fig. 4**a). Reconstructed images revealed abundant and densely arranged filaments on PSPMs derived from TIRAP-wt-expressing cells. In contrast, PSPMs from TIRAP-ΔPBM-expressing cells showed only sparse filamentous structures (**Fig. 4**b-d), indicating that any filaments formed by the mutant in intact cells were not stably bound to the PM upon isolation. Correspondingly, filament density was severely reduced for TIRAP-ΔPBM compared to TIRAP-wt (**Fig. 4**f). Averaging of multiple super-resolved filament segments revealed a helical, ribbon-like architecture with an average diameter of approximately 58 nm (**Fig. 4**g). Importantly, considering both the localization precision and linkage error intrinsic to DNA-PAINT^37,38^, the apparent dimension closely aligns with filament structures independently determined by cryo-EM (**Fig. 1**c). Averaging was not feasible for TIRAP-ΔPBM due to insufficient filament density on PSPMs.

### The PBM is essential for the functional coupling of TIRAP and MyD88 to TLR4

We next investigated whether the PBM as a structural requirement translates into functional consequences for TLR4 signaling. Receptor clustering is recognized as a critical mechanism for signal amplification in innate immunity, with studies demonstrating that spatial organization of TLR4 enhances MyD88-dependent signaling^39,40^. To quantitatively assess adaptor protein recruitment under spatially defined conditions that mimic receptor clustering, we devised a biofunctional nanodot array (bNDA) assay, which enables the analysis of protein-protein interactions in living cells^41,42^. The bNDA system utilizes nanopatterned surfaces fabricated by capillary nanostamping^43^ of PLL-PEG-ALFAtag, subsequently functionalized with an adaptor protein comprising anti-ALFAnb fused to tandem copies of the anti-mCherry DARPin 2m22 (aALFAnb-td2m22)^44^. Thus, tdmCherry-tagged TLR4 receptors in the plasma membrane of HeLa cells cultured on these substrates are selectively captured into bNDAs, enabling quantitative analysis of downstream effector recruitment (**Fig. 5**a).

**Fig. 5:**
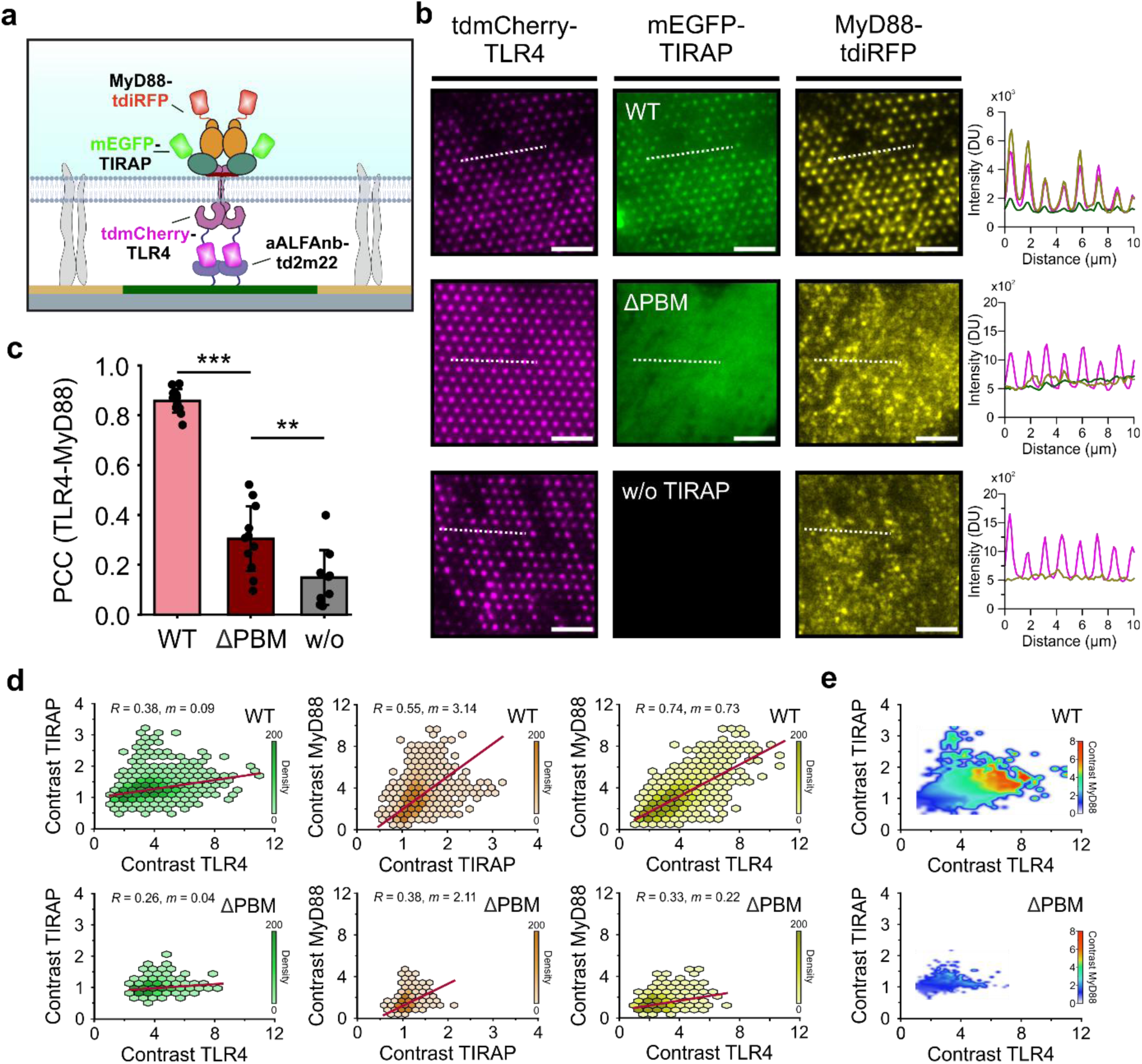
Functional coupling of TIRAP and MyD88 is disrupted by deletion of the PBM. **a** Schematic illustration of the biofunctional nanodot array (bNDA) assay. Nanopatterned surfaces were fabricated by capillary nanostamping of PLL-PEG-ALFAtag (green surface) and functionalized with anti-ALFAnb fused to tandem anti-mCherry DARPin 2m22 (aALFAnb-td2m22). TLR4 receptors tagged with tdmCherry (magenta) are selectively captured into bNDAs, facilitating recruitment of cytosolic mEGFP-TIRAP (green), which subsequently recruits MyD88-tdiRFP (orange). **b** Representative TIRF images of bNDAs showing tdmCherry-TLR4 (magenta), mEGFP-TIRAP (green), and MyD88-tdiRFP (yellow) for TIRAP-wt (top row), TIRAP-ΔPBM (middle row), and cells lacking TIRAP (bottom row). Line profiles show fluorescence intensity along the indicated dashed white lines. Scale bar: 5 µm. **c** Pearson correlation coefficients (PCC) between TLR4 and MyD88 fluorescence intensities per cell for TIRAP-wt (n = 11 cells), TIRAP-ΔPBM (n = 11 cells), and w/o TIRAP (n = 9 cells). Bar charts show mean ± s.d. with individual data points (***p < 0.001, **p < 0.01, one-way ANOVA with Tukey’s post hoc test). **d** Pairwise correlation analyses of contrast values for TLR4, TIRAP, and MyD88 are shown as hexbin density scatter plots for TIRAP-wt (top row) and TIRAP-ΔPBM (bottom row). Columns show TLR4-TIRAP (left), TIRAP-MyD88 (middle), and TLR4-MyD88 (right) correlations. Red lines indicate linear regression fits. Pearson correlation coefficients (*R*) and regression slopes (*m*) are indicated. Each analysis comprises > 4.000 individual nanodots from at least 9 cells per condition. **e** Two-dimensional density heatmaps showing MyD88 contrast as a function of TLR4 contrast (x-axis) and TIRAP contrast (y-axis) for TIRAP-wt (top) and TIRAP-ΔPBM (bottom).

We co-expressed tdmCherry-TLR4, mEGFP-TIRAP variants, and MyD88-tdiRFP as a functional readout and monitored protein recruitment at bNDAs by TIRF microscopy (**Fig. 5**b). Wild-type TIRAP efficiently colocalized with clustered TLR4 and robustly recruited MyD88, forming characteristic high-contrast protein interaction platforms (**Fig. 5**b, top row). In contrast, TIRAP-ΔPBM showed reduced recruitment to immobilized TLR4 and correspondingly diminished MyD88 accumulation (**Fig. 5**b, middle row). Control cells lacking TIRAP displayed successful TLR4 clustering but did not show MyD88 enrichment (**Fig. 5**b, bottom row; Supplementary Fig. 9). Pearson correlation analysis between TLR4 and MyD88 fluorescence intensities across individual cells confirmed these observations, revealing a strong positive correlation for TIRAP-wt (0.86), significantly weaker correlation for TIRAP-ΔPBM (0.30), and negligible correlation in the absence of TIRAP (0.15) (**Fig. 5**c).

To quantify adaptor and effector recruitment independently, we next analyzed mean fluorescence contrast values. Both TIRAP and MyD88 showed reduced co-recruitment in cells expressing TIRAP-ΔPBM compared to TIRAP-wt (Supplementary Fig. 10a, b). To account for variability in receptor density across bNDAs, we calculated relative MyD88-to-TLR4 contrast ratios, which confirmed significantly reduced coupling efficiency for TIRAP-ΔPBM, although partial function was retained above TIRAP-deficient controls (Supplementary Fig. 10c). This indicates that reduced MyD88 recruitment primarily reflects impaired TIRAP membrane accumulation rather than complete loss of intrinsic coupling capacity. Pairwise contrast-correlation analysis of thousands of individual nanodots revealed distinct assembly patterns (**Fig. 5**d). TIRAP-wt exhibited positive correlations across all three pairwise combinations, indicating tightly coordinated TIRAP accumulation and MyD88 recruitment at TLR4 clusters (**Fig. 5**d, top row). In contrast, TIRAP-ΔPBM displayed substantially weaker correlations with reduced coupling slopes (**Fig. 5**d, bottom row), while TIRAP-deficient controls showed no TLR4-MyD88 correlation (Supplementary Fig. 10d), confirming the absolute requirement for TIRAP in receptor-effector coupling. To visualize the functional interdependence of receptor clustering, adaptor recruitment, and effector engagement, we generated two-dimensional density heatmaps representing MyD88 contrast as a function of TLR4 and TIRAP levels (**Fig. 5**e). For TIRAP-wt, a distinct high-density region emerged at elevated TLR4 and TIRAP levels, suggesting that robust induction of MyD88 oligomerization requires a balanced ratio of co-clustered receptor and adaptor (**Fig. 5**e, top). This conclusion is further supported by the observation that MyD88 was much less efficiently recruited to highly clustered TIRAP at low TLR4 densities. In contrast, TIRAP-ΔPBM failed to generate such coordinated assembly patterns, instead showing a diffuse, low-density distribution characteristic of compromised signaling fidelity (**Fig. 5**e, bottom). Together, these analyses establish that the PBM is essential for steering TIRAP oligomerization at clustered TLR4 and subsequent MyD88 engagement, directly linking membrane binding capacity to downstream signaling competence.

## Discussion

Activation of TLR signaling complexes represents a fundamental example of how cells amplify external signals into switch-like downstream responses through cooperative protein interaction^45–47^. While TIR domain-containing adaptors have emerged as central players in achieving ultrasensitive signal propagation^5,27,28,33^, the molecular principles governing their nucleation and growth at membrane surfaces have remained poorly understood. Our integrated structural, biophysical, and cellular analysis of full-length human TIRAP reveals that the N-terminal PBM acts as a multifunctional regulatory module coupling membrane recognition to conformational activation and geometric constraint of filament assembly.

The 3.3 Å cryo-EM structure demonstrates that full-length TIRAP assembles into compact paired-strand ribbons rather than tubular architectures formed by isolated TIR domains^24^. This geometric difference is mechanistically significant: the ΔPBM variant retains its intrinsic assembly capacity, but forms wider, tubular assemblies and exhibits substantially reduced membrane affinity. The observation that the PBM remains unresolved in our reconstruction, combined with its predicted high disorder and the reported disorder-to-order transition upon lipid engagement^20^, suggests that this region regulates TIRAP oligomerization in addition to being a membrane anchor^48^. We propose that in the cytosolic environment, the flexible PBM maintains TIRAP in an assembly-incompetent, monomeric state. Such autoinhibitory function would explain why TIRAP TIR domains lacking the PBM form higher-order tubular aggregates more readily, and why cells can maintain a cytosolic pool of TIRAP without constitutive activation of inflammatory signaling. Critically, the sequential oligomerization pathway establishes nucleation as the rate-limiting step, providing an additional kinetic barrier against spontaneous cytosolic aggregation^49^. Spontaneous assembly of full-length TIRAP into more condensed filaments, which are probably not capable to induce Myddosome assembly, may point to another tier to restrain constitutive signal activation.

Our reconstitution experiments on supported lipid bilayers reveal that membrane composition profoundly controls not only recruitment, but also the assembly mechanism. On neutral DOPC membranes, TIRAP binding follows single-exponential kinetics characteristic of simple adsorption. In contrast, PIP_2_-containing bilayers induce sigmoidal kinetics with pronounced lag phases followed by accelerated cooperative growth. These observations indicate that PIP_2_ not only enhances membrane recruitment via electrostatic interactions with PBM^19–21^ but surprisingly also inhibits oligomerization, which would be expected to be accelerated due to the high local concentration. Although high TIRAP concentrations induced spontaneous filament assembly both *in vitro* and in cells, we assume that at physiological concentrations, which are substantially lower, PBM-mediated membrane binding likely maintains TIRAP in a monomeric state. However, the observed kinetics indicate that once oligomerization is initiated, cooperative growth proceeds rapidly. Using DNA-PAINT super-resolution microscopy, we provide the first direct ultrastructural visualization of TIRAP filaments at cellular plasma membranes, which closely resemble the paired-strand ribbon architecture observed *in vitro*, i.e. the postulated inactive conformation.

Live-cell nanodot array experiments identified that clustered TLR4 receptors efficiently recruit full-length TIRAP, which in turn initiates accumulation of MyD88 as expected for propagating downstream signaling. Interestingly, lack of PBM abrogated binding of TIRAP to TLR4, establishing a direct link between membrane retention capacity and signaling competence. Analysis of individual nanodots revealed that productive complex formation requires coordinated accumulation of both receptor and adaptor, with MyD88 recruitment being maximal only when TLR4 and TIRAP are both present at elevated local densities. These observations argue against a model, in which pre-formed TIRAP filaments engage receptors, and instead support receptor clustering as the initiating event triggering TIRAP oligomerization.

Based on our findings, we propose a four-phase model for membrane-gated TIRAP assembly (**Fig. 6**). In the resting state, TIRAP exists predominantly in an assembly-incompetent state, with the flexible PBM preventing productive oligomerization. TIRAP dynamically associates with the plasma membrane through PBM-mediated interactions, and the intrinsically disordered PBM adopts helical structure upon membrane engagement. Anchoring TIRAP to negatively charged lipids at the plasma membrane maintains the TIR domains in a conformation/orientation likely incompatible with oligomerization. Ligand-induced TLR4 dimerization triggers spatial reorganization by recruiting TIRAP to activated receptor complexes through TIR-TIR interactions, thus locally enriching and scaffolding productive TIRAP oligomerization. The resulting membrane-anchored TIRAP filament in turn templates MyD88 recruitment via heterotypic TIR-TIR interactions, with the filament geometry potentially defining the stoichiometry required for efficient Myddosome assembly^17,18,50,51^. This mechanism exemplifies the SCAF paradigm^28^ wherein ultrasensitivity emerges from cooperative assembly rather than enzymatic cascades, transforming graded receptor occupancy into threshold-dependent signal propagation.

**Fig. 6:**
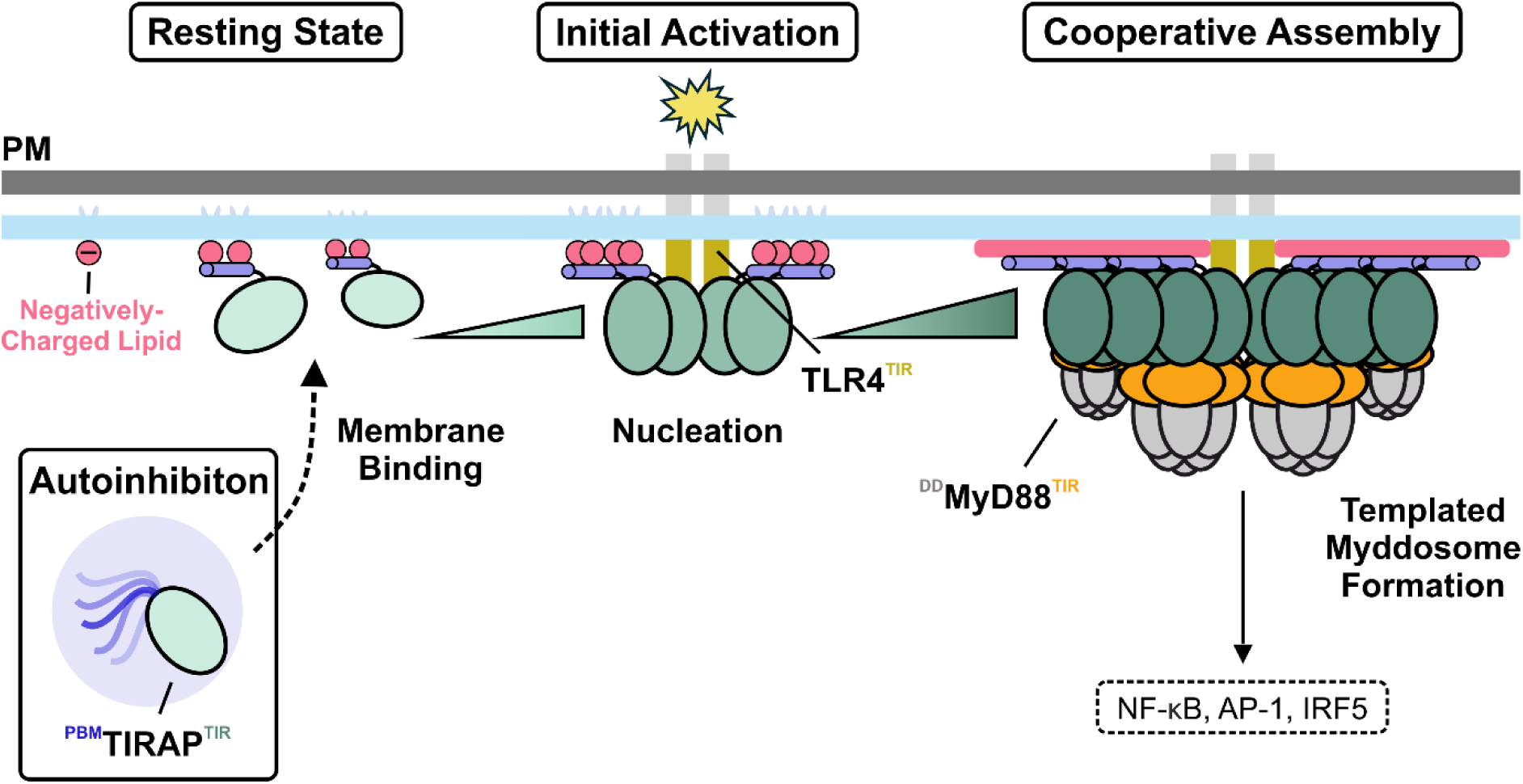
Four-phase model for membrane-gated TIRAP assembly. Schematic illustration depicting the mechanistic progression from resting state to productive signaling complex formation. In the resting state, TIRAP (green, TIR domain; blue, PBM) exists in an assembly-incompetent conformation with the flexible PBM preventing productive oligomerization. TIRAP dynamically associates with the plasma membrane via PBM-mediated interactions, maintaining a conformation incompatible with oligomerization. Initial Activation: Ligand-induced TLR4 dimerization recruits TIRAP via its TIR domains (yellow), thus initiating nucleation of oligomers. Cooperative Assembly: When local concentration exceeds the critical threshold, cooperative nucleation initiates filament growth. TIRAP filaments template MyD88 (grey, death-domains DD; orange, TIR domain) recruitment forming Myddosomes that activate downstream signaling.

In conclusion, our study establishes TIRAP as a membrane-organized molecular switch. The PBM serves multiple regulatory functions by gating assembly competence through membrane engagement, constraining filament geometry compatible with effector recruitment, and enabling stable signaling platform formation at sites of receptor activation. These findings provide a comprehensive structure-function framework for TIRAP and reveal general principles by which membrane-organized cooperative assembly converts graded receptor occupancy into threshold-dependent signal propagation.

## Methods

### Purification of human TIRAP from *E. coli*

Human TIRAP (UniProt: P58753) was synthesized by Genescript and subcloned into pET28a vector, incorporating a TEV protease-cleavable N-terminal His_6_-Myc tag. The construct was transformed into *Escherichia coli* BL21 (DE3) Rosetta cells. For protein expression, cells were cultured at 37°C in Lysogeny Broth (LB) medium to an optical density (OD_600_) of 0.6. Protein expression was induced with 0.2 mM isopropyl β-D-1-thiogalactopyranoside (IPTG) and cells were incubated for 20 h at 20°C. Cells were harvested by centrifugation (4000 × *g*, 15 min, 4 °C), and resuspended in lysis buffer containing 50 mM HEPES pH 7.5, 500 mM NaCl, 1 mM dithiothreitol (DTT), 1 mM phenylmethylsulfonyl fluoride (PMSF), 10 µg/ml DNase I, and 5 mM MgCl_2_. Cells were lysed by sonication (Branson Ultrasonics, Brookfield, USA), and the lysate was clarified by centrifugation at 40,000 × *g* for 30 min at 4°C. After an incubation for 2 h on a nutator with pre-equilibrated His-Pur Ni-NTA Sepharose (Thermo Fisher Scientific), beads were washed thoroughly, and TIRAP was eluted in elution buffer (50 mM HEPES, pH 7.5, 150 mM NaCl and 1 mM DTT) containing 250 mM Imidazole. Pooled samples were desalted using a PD MiniTrap G-25 desalting column (Cytiva) and further purification was performed by anion-exchange chromatography using a 1 ml HiTrap Q HP column (Cytiva) on an ÄKTA pure FPLC system (Cytiva, Marlborough, USA). hTIRAP was eluted at 400 mM NaCl using an isocratic gradient from buffer A (20 mM Tris-HCL pH 8.0) to buffer B (20 mM Tris-HCL pH 8.0, 1 M NaCl). Protein purity was confirmed by SDS-PAGE with Coomassie Brilliant Blue staining (Supplementary Fig. 2b). Purified TIRAP was aliquoted, frozen in liquid nitrogen, and stored at −80°C. The phosphomimetic variant T28D and N-terminal deletion mutant Δ79 (residues 80-221) were generated by site-directed mutagenesis and PCR-based deletion mutagenesis, respectively, using the Gibson Assembly Master Mix (New England Biolabs, #M5510). Proteins were expressed and purified following the same protocol as above mentioned.

### Negative stain electron microscopy

Protein sample quality was examined by negative-stain electron microscopy using 2 % (w/v) uranyl formate solution as previously described^52^. Negative-stain micrographs were collected manually on a JEM-2100Plus transmission electron microscope (JEOL) operated at 200 kV and equipped with a XAROSA CMOS 20-megapixel camera (EMSIS) at a nominal magnification of 30,000 (3.12 Å per pixel). Data shown in Supplementary Fig. 2c, d.

### Cryo-EM sample preparation and data acquisition

For cryo-EM studies, 0.9 mg/ml TIRAP was incubated for 1 h at 37°C to induce filament formation. C-flat grids (CF-1.2/1.3-3Cu-50, Protochips Inc., USA) were glow-discharged using a PELCO easiGlow device at two consecutive runs of 15 mA for 45 s. A sample volume of 3 µL was applied to the grid and immediately plunge frozen in liquid ethane using a Vitrobot Mark IV (Thermo Fisher Scientific) with the environmental chamber set at 100 % humidity and 20°C. All datasets were collected on a Glacios cryo-transmission electron microscope (Thermo Fisher Scientific) operated at 200 kV and equipped with a Selectris energy filter (Thermo Fisher Scientific) set to a slit width of 10 eV. Movies were recorded using a Falcon 4 direct electron detector (Thermo Fisher Scientific) at a nominal magnification of 130,000, corresponding to a calibrated pixel size of 0.945 Å per pixel. Movies were aquired in the Electron-Event Representation (EER) format and recorded at a total dose of of 50 e⁻/Å². Automated data collection was performed using EPU software version 2.9 (Thermo Fisher Scientific) with a defocus range of −0.8 to −2.0 μm.

### Cryo-EM data processing

All cryo-EM datasets were processed using cryoSPARC (v.4.0)^53^. The complete processing workflows are shown in Supplementary Fig. 3. Movies were preprocessed with patch-based motion correction, patch-based contrast transfer function (CTF) estimation and filtered by CTF fit estimates using a 4.5 Å cutoff.

Filament positions were determined using the filament tracer tool with an expected filament diameter of 30-100 Å. Picked positions were manually curated using curvature < 0.001 Å^−1^ and sinuosity < 1 cutoffs. A stack of 2,211,683 particles were subjected to two rounds of 2D classification, which resulted in 785,751 particles corresponding to well-aligning 2D projections. Initial helical refinement with non-uniform regularization during the refinement without supplied helical symmetry parameters resulted in a 3.8 Å reconstruction. Helical symmetry parameters were determined using the helical symmetry search tool in cryoSPARC, yielding a helical rise of 16.6 Å and a helical twist of −176.02°. A final helical reconstruction with supplied helical symmetry parameters resulted in a global resolution of 3.3 Å.

Details about data collection for individual datasets and validation statistics are given in Supplementary Table 1. Reported B-factors resulted from unsupervised auto-sharpening during refinement in cryoSPARC. To aid model building, unsharpened half-maps, FSC-based resolution and the molecular mass were supplied as input for density modification within Phenix^54^ (*phenix.resolve_cryo_em*).

### Model building, refinement and validation

Initial atomic models were generated using the AlphaFold2 prediction of human TIRAP (UniProt: P58753). The predicted structure was rigid-body fitted into the cryo-EM density map using UCSF ChimeraX^55^. Manual model building and adjustment were performed in Coot v0.9^56^ and iterative real-space refinement was performed using *phenix.real_space_refine* within Phenix^54^. Validation reports were automatically generated by MolProbity^57^ within Phenix. The final atomic coordinates and corresponding cryo-EM density map have been deposited in the Protein Data Bank (PDB ID: 9FQM) and the Electron Microscopy Data Bank (EMDB ID: EMD-50657).

### Gradient fixation

Gradient fixation was performed as previously reported^30^. Briefly, 0.9 mg/ml TIRAP was incubated at 37°C for 1 h to induce filament formation. Large aggregates were dissolved by brief sonication. A continuous glycerol-glutaraldehyde gradient was prepared using a Gradient Master (BioComp Instruments, Fredericton, Canada). The light solution (top layer) consisted of 50 mM HEPES pH 7.5, 200 mM NaCl, and 15 % (v/v) glycerol. The heavy solution (bottom layer) contained 50 mM HEPES pH 7.5, 200 mM NaCl, 45 % (v/v) glycerol, and 0.1 % (v/v) glutaraldehyde. Gradients were pre-incubated at 4°C for 1 h to allow equilibration and initiate crosslinking. Subsequently, 180 pmol of TIRAP was carefully added to the gradient and crosslinked at 22,500 × *g* for 14 h at 10°C. Following centrifugation, desired fractions were analyzed by SDS-PAGE.

### Mass photometry

Mass photometry experiments were performed to monitor TIRAP oligomerization kinetics in solution using a Refeyn TwoMP mass photometer (Refeyn Ltd., Oxford, UK). The instrument was calibrated using native protein standard (β-amylase) according to the manufacturer’s instructions. Data were acquired using AcquireMP software and analyzed using DiscoverMP v2023 (both Refeyn Ltd) by fitting with Gaussians and plotted as normalized counts. High Precision Cover Glasses (Marienfeld) were used for sample analysis. Perforated silicone gaskets were placed on the coverslips to form wells for every sample to be measured. For non-oligomerizing control conditions, TIRAP was diluted to 100 nM on ice for 1 h prior to measurement. To monitor oligomerization, 500 nM TIRAP was analyzed after incubation for 0, 15 and 30 min at 37°C.

### Lipid protein overlay assay

Lipid-binding specificity of TIRAP was assessed using an established protein-lipid overlay assay^58^. Briefly, 10 µg of desired lipids (Anatrace) were manually spotted onto Immobilon-P polyvinylidene difluoride (PVDF, Merck Millipore, IPVH00010) membranes, dried for 1 h at room temperature and blocked with PBS-T (0.1 % (v/v) Tween-20; 3 % (w/v) fatty acid-free bovine serum albumin (BSA)) for 1 h at room temperature. Membranes were then incubated with 5 µg/ml His-tagged TIRAP (wild-type, T28D, or Δ79 variant) ) for 2 h at room temperature.

Following incubation, membranes were washed with PBS-T and incubated with monoclonal anti-His antibody (Takara Bio, Cat. 631212, 1:5000 dilution) in blocking buffer for 1 h at room temperature. Membranes were washed with PBS-T and incubated with horseradish peroxidase (HRP)-conjugated goat anti-mouse IgG secondary antibody (Santa Cruz Biotechnology, Cat. sc-516102; 1:1000 dilution) in blocking buffer for 1 h at room temperature. Membranes were washed and bound TIRAP was detected using an ECL detection kit (SuperSignal West Pico PLUS, Cat. 34577).

### Lipid preparation and supported lipid bilayer formation

Lipid were purchased from commercial sources: 1,2-dioleoyl-*sn*-glycero-3-phosphocholine (DOPC; Avanti Polar Lipids, Cat. 850375), L-α-phosphatidylinositol-4,5-bisphosphate (PI(4,5)P_2_; Avanti Polar Lipids, Cat. 840046), and Oregon Green™ 488 1,2-dihexadecanoyl-*sn*-glycero-3-phosphoethanolamine (OG488-DHPE; Thermo Fisher Scientific, Cat. O12650). Lipids were mixed in chloroform at the desired molar (%) ratios (94:5:1 DOPC:PIP_2_:OG488-DHPE or 99:1 DOPC:OG488-DHPE), evaporated under nitrogen flux, and dried under vacuum overnight. The resulting lipid films were rehydrated in HEPES-buffered saline (HBS: 20 mM HEPES pH 7.4, 150 mM NaCl) to a final lipid concentration of 0.7 mg/ml and subjected to ten freeze-thaw cycles. Small unilamellar vesicles (SUVs) were generated by manual extrusion using an Avanti Mini-Extruder (Avanti Polar Lipids, Cat. 610020), first by 11 passages through polycarbonate membranes with 100 nm pore size (Sigma-Aldrich, Cat. WHA110605), followed by 15 passages through 50 nm pore-size membranes (Sigma-Aldrich, Cat. WHA110404).

For supported lipid bilayer (SLB) formation, glass coverslips (Ibidi µ-Slide 8 Well, Cat. 10812, #1.5H) were cleaned using piranha solution to ensure complete removal of organic contaminants. Piranha solution was freshly prepared by carefully adding hydrogen peroxide (Sigma-Aldrich, Cat. 95294) to concentrated sulfuric acid (Carl Roth, Cat. X942.1) in a 1:3 (v/v) ratio in a glass beaker. Coverslips were immersed in piranha solution for 60 min at room temperature, rinsed with deionized water for 5 min and dried under a nitrogen stream. Coverslips were further cleaned by plasma treatment (Plasma Cleaner Femto 1A, Diener Electronic GmbH) for 2 min immediately before usage and were mounted onto µ-Slide sticky-Slide 8 Well (Ibidi, Cat. 80828). SUVs were incubated at a final lipid concentration of 0.2 mg/ml at 37°C for 10 min with 3 mM CaCl_2_ and washed 15 times with HBS buffer to remove non-fused vesicles.

### Fluorescence recovery after photobleaching

Bilayer integrity and lateral lipid mobility were confirmed by fluorescence recovery after photobleaching (FRAP) of the fluorescent lipid tracer OG488-DHPE. Three pre-bleach TIRF images were acquired, followed by photobleaching of a circular region of interest (ROI, 10 µm diameter). Fluorescence recovery was monitored by acquiring 100 frames at 4 frames per second (250 ms per frame). Mean fluorescence intensity within the bleached ROI was quantified using Fiji^59^. Recovery curves were corrected for overall photobleaching by normalizing to an unbleached reference ROI and normalized to the maximum pre-bleach intensity using OriginPro9.

### Time-lapse imaging and analysis of TIRAP on supported lipid bilayers

For visualization of TIRAP on SLBs, recombinant His_6_-TIRAP was fluorescently labeled by pre-incubation with Dy547-trisNTA loaded with Ni^2^ ions^31,32^ at a final molar ratio of 10:1 (protein:dye) for 3 min on ice, and then added to SLBs of defined lipid composition (working concentrations 1 µM or 10 µM). Real-time imaging of TIRAP binding and filament formation on SLBs was performed by total internal reflection fluorescence (TIRF) microscopy at 37°C using the Olympus IX-83 TIRF system described below. For each condition, time-lapse movies were recorded at a frame rate of 1 frame per second (fps) for a total duration of 400 s using 561 nm excitation and the ORCA-FusionBT sCMOS camera with 2×2 binning (effective pixel size: 130 nm). Camera exposure time was set to 30 ms per frame. To minimize photobleaching, laser power was optimized to maintain signal-to-noise ratio while preventing excessive fluorophore bleaching. Two independent samples were analyzed per condition, with multiple regions of interest (ROIs; n = 12 per condition) imaged per experiment. For each ROI (13 × 13 µm), mean fluorescence intensity over time was quantified using Fiji^59^. Background fluorescence was subtracted from all measurements. Fluorescence intensity traces were normalized to the first frame (t = 0 s) to calculate relative intensity (*I_rel_*) over time.

Normalized intensity traces were analyzed to extract quantitative assembly parameters using custom Python scripts. For DOPC bilayers, TIRAP binding followed simple exponential kinetics and was fitted with a single-exponential function:

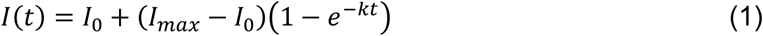

where *I_0_* is the baseline fluorescence intensity at *t* = 0, *I*_max_ is the plateau intensity, *k* is the exponential rate constant, and *t_1/2_* = ln(2)/*k* is the half-time.

For PIP_2_-containing bilayers, TIRAP assembly exhibited sigmoidal kinetics characteristic of cooperative assembly and was fitted using the Finke-Watzky^60^ two-step nucleation-growth model. The model describes the time-dependent concentration of assembled TIRAP (*B(t)*) as:

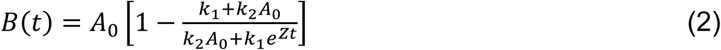

where *A_0_* is the initial TIRAP concentration (normalized to maximum signal), *k_1_* is the nucleation rate constant, *k_2_* is the autocatalytic growth rate constant, and *Z* = *k_1_* + *k_2_A*_0_. From the fitted parameters, the following kinetic descriptors were calculated:

- Half-time (*t_1/2_*): Time to reach 50% of plateau intensity, calculated as *t*_1/2_ = ln(*k*₂*A*₀/*k*₁)/*Z*
- Time of maximum slope (*t_max_*): Time at which assembly rate is maximal, *t_max_* = ln(*k*₂*A*₀/*k*₁)/*Z*
- Maximum assembly slope (*m_max_*): Maximum first derivative of the fitted curve, *m_max_*=(*k*₁+*k*₂*A*₀)²/(4*k*₂)
- Induction time (*t_ind_*): Time corresponding to the end of the lag phase, calculated using the jerk method as *t_ind_* = ln[(2−√3)×*k*₂*A*₀/*k*₁]/*Z*

All curve fitting was performed using the *scipy.optimize.curve_fit* function. Initial parameter estimates for *k_1_* and *k_2_* were manually defined based on visual inspection of the kinetic traces to ensure convergence. Goodness of fit was assessed by calculating the coefficient of determination (*R*²). All fits exhibited *R*² > 0.98. Detailed kinetic parameters for all experimental conditions are provided in Supplementary Table 2.

To quantify the spatial extent of TIRAP assembly on SLBs, time-lapse movies were processed in Fiji^59^ to segment filamentous structures and calculate membrane coverage. Raw fluorescence images were first subjected to a 2-pixel top-hat filter to enhance filamentous structures by removing uneven background. Subsequently, rolling-ball background subtraction was applied with a radius of 5 pixels. Afterwards, filamentous structures were segmented using the Trainable Weka Segmentation v3.3.4 plugin^61^ in Fiji. A binary classifier was trained on representative image stacks to distinguish two classes: (1) filamentous structures and (2) non-filamentous background. Training was performed interactively by manually annotating filaments and background regions in multiple frames. After training, probability maps corresponding to the class (1) were generated for all frames. Probability maps were converted to binary masks using the automatic minimum threshold method. Segmented binary stacks were skeletonized to extract filament centerlines. For temporal visualization, skeletonized stacks were split into subsets and Z-projected using sum intensity to generate time-binned images representing cumulative filament density over the acquisition period.

Membrane coverage by TIRAP filaments was calculated from endpoint images in Fiji. Multiple ROIs (n > 25 ROIs per condition across 2 independent experiments) were analyzed. Each ROI was processed with a 2-pixel top-hat filter to enhance contrast. Binary segmentation was performed using an automatic threshold (Default method). The membrane area covered by TIRAP was quantified using the Particle Analyzer tool.

### Cell culture and transfection

HeLa cells were cultured in Minimal Essential Medium (MEM) with Earle’s salts and phenol red (PanBiotech, Cat. P04-09500) supplemented with 10 % (v/v) fetal bovine serum (FBS; PanBiotech, Cat. P30-3031), 2 mM L-alanyl-L-glutamine, 1 % (v/v) non-essential amino acids (MEM NEAA, PanBiotech, Cat. P08-32100), and 10 mM HEPES buffer (PanBiotech, Cat. P05-01100). Cells were cultured in a humidified incubator at 37°C with 5 % CO_2_.

For transient transfection, HeLa cells were seeded into 60 mm tissue culture dishes and incubated overnight to achieve 70-80 % confluence at the time of transfection. Transfection was performed using Lipofectamine™ 2000 Transfection Reagent (Thermo Fisher Scientific, Cat. 11668027) according to the manufacturer’s protocol with minor modifications. For each 60 mm dish, 5 µg of plasmid DNA (see Supplementary Table S3 for plasmid details) was diluted in 300 µl of Opti-MEM™ Reduced Serum Medium (Thermo Fisher Scientific, Cat. 31985070) in a sterile 1.5 ml microcentrifuge tube and mixed gently by pipetting. In a separate tube, 10 µl of Lipofectamine™ 2000 reagent was diluted in 300 µl of Opti-MEM™ and mixed gently. Both solutions were incubated separately for 5 min at room temperature to allow equilibration. Afterwards, both solutions were combined, mixed gently, and incubated for 20 min at room temperature. The transfection mixture was then added dropwise to the culture dishes. After 4-6 h incubation, the medium was replaced with 4 ml of fresh pre-warmed complete MEM. Cells were incubated for an additional 24 h to allow protein expression.

For microscopy experiments, transfected cells were detached by Trypsin/EDTA (PanBiotech, Cat. P10-024100) treatment and seeded on microscopy glass coverslips (Marienfeld Laboratory Glassware, Cat. 0117640). Coverslips were first cleaned by immersion in isopropanol, followed by rinsing with deionized water. Coverslips were dried under a stream of nitrogen gas and subsequently cleaned by plasma treatment (Plasma Cleaner Femto 1A, Diener Electronic GmbH) at 100 % power for 15 min. Subsequently, coverslips were functionalized with poly-L-lysine coupled to a polyethylene glycol functionalized with RGD-peptide (PLL-PEG-RGD)^62^ to facilitate cell attachment. Coverslips were then rinsed three times with PBS (PanBiotech, Cat. P04-36500) to remove unbound polymer and equilibrated with 500 µl complete MEM. For experiments requiring alternative surface functionalization (e.g., polymer-supported plasma membranes or biofunctional nanodot arrays), specific coating protocols are described in the respective sections below.

### Total internal reflection fluorescence microscopy (TIRFM)

TIRF imaging was performed at 25°C (37°C for *in vitro* experiments on SLBs) using an inverted microscope (Olympus IX-83) equipped with a modular two-deck configuration. The upper deck was equipped with a 4-line TIRF condenser (cellTIRF (MITICO), Olympus), while the lower deck contained the cellFRAP module (Olympus) for photobleaching experiments. The system was equipped with multiple laser excitation sources, including a 405 nm laser (BCL-100-405, CrystaLaser), a 488 nm diode laser (LuxX 488-200, Omicron), a 561 nm fiber laser (2RU-VFL-P-500-561-B1R, MPB Communications) and a 642 nm fiber laser (2RU-VFL-P-500-642-B1R, MPB Communications). Fluorescence excitation and emission were controlled via a TIRF pentaband polychroic beamsplitter (Semrock zt405/488/561/640/730rpc) and a penta-bandpass emitter filter (BrightLine HC 440/521/607/694/809). Additional single-bandpass filters were used for channel-specific detection: blue (Semrock BrightLine HC 445/45), green (Semrock BrightLine HC 525/35), orange (Chroma 600/50 ET) and red (Chroma 685/50). The system was further equipped with a motorized ultrasonic XY-stage (IX3-SSU, Olympus), and a hardware autofocus system (IX3-ZDC2, 830 nm version, Olympus). Images were acquired with a Hamamatsu ORCA-FusionBT sCMOS camera (2304×2304 pixel resolution) using a 100× oil immersion objective (UPLAPO 100× HR, NA 1.5, Olympus). Imaging was controlled via CellSens Dimension software version 3.1 (Olympus).

For FRAP experiments, the cellFRAP module (Olympus) was utilized with a 405 nm bleaching laser (LuxX+ 405-60, Omicron). Image acquisition was conducted using a three-line polychroic beamsplitter (Chroma, zt488/561/633/405tpc) located in the upper deck, while the lower deck incorporated a FRAP cube consisting of a dichroic beamsplitter at 405 nm (Chroma, H 405 LPXR superflat) and a QuadLine emitter filter (Chroma, zet405/488/561/640m).

### DNA-PAINT imaging, processing and analysis

Samples for DNA-PAINT imaging were fixed in PBS containing 3 % (w/v) paraformaldehyde and 0.1 % (v/v) glutaraldehyde for 15 min at room temperature, washed five times with PBS, permeabilized with 0.1 % (v/v) Triton-X100 (Sigma-Aldrich, Cat. T8787) for 10 min, and blocked with 3 % (w/v) BSA in PBS for 30 min. Samples were incubated overnight at 4°C with 50 nM anti-GFP nanobody conjugated to docking strand (MASSIVE-TAG-X2-FAST anti-GFP, Massive Photonics), washed three times with PBS, and post-fixed with 3 % paraformaldehyde and 0.1 % glutaraldehyde for 15 min. After washing, 90 nm gold nanoparticles (Cytodiagnostics, Cat. G-90-20, 1:1000 dilution) were added as fiducial markers for 15 min, followed by additional washing steps.

DNA-PAINT imaging was performed at 25°C using the previously described TIRF microscopy setup. Samples were imaged in imaging buffer containing 100 pM Cy3B-conjugated imager strand F3 (MASSIVE-TAG-X2-FAST anti-GFP, Massive Photonics), with 561 nm excitation at approximately 100 W cm^−^² power density in the focal plane, which was consistent for all experiments. Datasets of 40000 frames were acquired at 50 ms exposure time with 2×2 pixel binning (effective pixel size: 130 nm). A hardware autofocus system ensured stable focus throughout the acquisition.

Raw datasets were processed using the Picasso (version 0.7.5) software suite^34^. Single-molecule localizations were identified using Picasso Localize (box size: 7 pixels; minimum net gradient: 5000-10,000) using the following photon conversion parameters: EM gain: 1; baseline: 400; sensitivity: 0.26; quantum efficiency: 0.92; pixel-size: 130 nm. Localizations were fitted with an integrated Gaussian function^63^, filtered for localization precision ≤ 6.5 nm (0.05 pixels) and drift-corrected using cross-correlation^64^ refined with fiducial markers. Averaged reconstruction images were generated using Picasso Average by aligning 50-100 filament fragments (oversampling: 40; iterations: 10). Filament width and length were measured manually from reconstructions. For filament density quantification, localizations were exported to Fiji^59^, processed with a tubeness filter, and segmented using Trainable Weka Segmentation v3.3.4^61^ with two classes (filamentous structures and background). Probability maps were thresholded using Otsu’s method, refined with watershed segmentation, and segments < 10 pixels were excluded.

### Confocal laser-scanning microscopy (cLSM)

Confocal images were acquired with an inverted Olympus IX83-P2ZF microscope equipped with a motorized 8-position filter cassette (FV30-RFACA, Olympus), motorized resolving nosepiece, motorized xy-stage (IX3-SSU, Olympus), xy-piezostage for confocal point scanning (P-542.2SL, Physik Instrumente), a condenser IX2-LWUCD (working distance 27 mm, NA 0.55) and an UPLSAPO 60x water immersion objective (NA 1.2, Olympus). Emitted photons were detected using photomultiplier tubes (Hamamatsu R7862). Coarse adjustment was performed with an epifluorescence mercury lamp (100 W, Olympus). For each scan, the dichroic beam splitter DM 405/488/550/635 was used. Excitation of mEGFP (488 nm) was performed with an argon laser (Showa Optronics Argon Multiline Laser 458/488/515 GLG3135) at 0.1-1 % laser power depending on expression level. Zoom was set to 4× o capture single cells and scanning speed was set to 40 µs per pixel. Fluorescence intensity ratios were calculated by dividing plasma membrane fluorescence by cytosolic signal using Fiji^59^.

### Preparation of polymer-supported plasma membranes (PSPMs)

PSPMs were generated according to previously published protocols^36,65^. Briefly, microscopy glass coverslips (Marienfeld Laboratory Glassware, Cat. 0117640) were immersed in isopropanol, dried and further cleaned by plasma cleaning (Plasma Cleaner femto 1A, Diener electronics) at 100 % power for 15 min. Coverslips were coated with a 70:30 (w/w) mixture of PLL-PEG-HTL and PLL-PEG-RGD^62^. Transfected HeLa cells co-expressing mEGFP-TIRAP (wild-type or ΔPBM) and HaloTag-mTagBFP-TMD were seeded onto functionalized coverslips one day prior to PSPM generation. The RGD-peptide functionalization enables adherence of the cells by integrin interactions, while the covalent HTL-HaloTag interaction allows tethering of the plasma membrane to the surface. PSPMs were produced by incubating cells with 10 µM Latrunculin B (Abcam, Cat. Ab144291) for 5 min at 37°C, followed by removal of the cell body by sheer forces through heavy pipetting. Immediately afterward, PSPMs were fixed using 3% (w/v) paraformaldehyde and 0.1% (v/v) glutaraldehyde in PBS for 15 min at room temperature.

### Biofunctional nanodot arrays (bNDAs) and interaction analysis

Surface modification, synthesis of mesoporous silica stamps, and preparation of patterned substrates were conducted following previously established protocols^41–43,66^. Briefly, glass coverslips (Marienfeld Laboratory Glassware, Cat. 0117640) were plasma-cleaned (Plasma Cleaner Femto 1A, Diener Electronic GmbH), silanized with vinyltrimethoxysilane (VTMS; Sigma-Aldrich, Cat. 235768) and subsequently nanopatterned by capillary nanostamping of PLL-PEG-ALFAtag. After stamping, substrates were backfilled with a 90:10 (v/v) mixture of PLL-PEG-OMe and PLL-PEG-RGD^62^ at 1.0 mg ml^−1^ for 30 min, rinsed thoroughly with ultrapure water, and stored at 4°C. Immediately prior to cell seeding, substrates were incubated with 1 µM anti-ALFAnb-tandem-2m22DARPin adapter protein for 30 min to enable selective immobilization of tdmCherry-tagged TLR4 receptors. HeLa cells co-expressing tdmCherry-TLR4, mEGFP-TIRAP (wild-type or ΔPBM), and MyD88-tdiRFP were seeded onto functionalized substrates and incubated 4-6 hours. Cells were imaged by TIRF microscopy as described above. Multi-channel image stacks were acquired with identical exposure settings within each experimental condition.

Image analysis was performed using a custom MATLAB script (MathWorks, version R2023b). Individual channels were loaded and calibrated using pixel size (130 nm) and camera offset (400). Analysis regions were manually defined by drawing polygonal masks on the bait channel (TLR4) to exclude imaging artifacts, surface defects, or regions with non-specific binding. Individual nanodots were detected using Laplacian of Gaussian (LoG) filtering with automatic threshold determination. Local minima in the LoG-filtered image below threshold were identified as potential bNDAs and filtered based on roundness and minimum center-to-center distance to eliminate false positives and overlapping events. For each detected bNDA, annular background regions were automatically generated around the spot periphery, ensuring local background sampling while avoiding neighboring spots. Regions with insufficient area coverage were excluded. Local background intensities were calculated by expanding ring masks by one additional pixel and quantifying mean pixel intensities within these extended annular regions, restricted to predefined analysis regions. For each nanodot, dot intensity (*I_dot_*), local background intensity (*I_bg_*), and contrast *C* were calculated as:

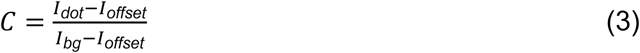

where *I_offset_* represents the camera offset. Prey channel images (TIRAP, MyD88) were analyzed at identical nanodot positions. To account for variability in receptor density, relative contrast values were determined by normalizing prey contrast to corresponding bait contrast:

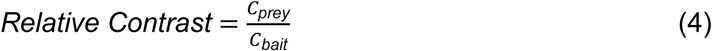

Distributions of contrast and relative contrast values were visualized as box plots. Statistical significance between experimental conditions (TIRAP-wt, TIRAP-ΔPBM, and TIRAP-deficient controls) was assessed using two-sample Kolmogorov-Smirnov tests. Pairwise correlations between TLR4, TIRAP, and MyD88 intensity and contrast values were calculated using Pearson correlation coefficients. To visualize relationships between receptor clustering and adaptor recruitment, hexbin scatter plots and two-dimensional density heatmaps were generated using Python version 3.13.

## Data availability

All data generated or analyzed during this study are included in the manuscript and supporting file. All density maps and models have been deposited in the Protein Data Bank (PDB) under ID 9FQM and the Electron Microscopy Data Bank (EMDB) under ID EMD-50657. Structures were visualized with ChimeraX^55^ and protein interactions were analyzed with the help of PDBe PISA^67^ and LigPlot^68^. An overview of the local density fit is given in Supplementary Fig. 3e.

## Acknowledgements

We thank A. Budke-Gieseking, G. Hikade, H. Kenneweg, and W. Kohl for excellent technical support. We are grateful to members of the Piehler and Möller laboratories for helpful discussions. J.H.S was supported by the Friedrich-Ebert Foundation. Funding: this work was supported by the Deutsche Forschungsgemeinschaft (DFG, German Research Foundation, SFB 1557, 467522186) to A. M. and J.P.

## Author contributions

A.F., J.H.S., K.T. and M.W. performed all experiments and analyzed data. A.F., J.H.S., A.M. and J.P. conceptualized the project. A.F. and J.H.S. wrote the original manuscript draft with input from A.M. and J.P. All authors contributed to manuscript revision and approved the final version. A.M. and J.P. secured funding for this study.

## References

1 Akira, S., Uematsu, S. & Takeuchi, O. Pathogen Recognition and Innate Immunity. Cell 124, 783–801 (2006). 10.1016/j.cell.2006.02.015

2 Chen, R., Zou, J., Chen, J., Zhong, X., Kang, R. & Tang, D. Pattern recognition receptors: function, regulation and therapeutic potential. Signal Transduction and Targeted Therapy 10, 216 (2025). 10.1038/s41392-025-02264-1

3 Chen, Y.-H., Wu, K.-H. & Wu, H.-P. Unraveling the Complexities of Toll-like Receptors: From Molecular Mechanisms to Clinical Applications. International Journal of Molecular Sciences 25, 5037 (2024).

4 Alexopoulou, L. & Irla, M. Toll-like receptors (TLRs) in the trained immunity era. eLife 14, e106443 (2025). 10.7554/eLife.106443

5 Rughetti, A. et al. Imperative role of adaptor proteins in macrophage toll-like receptor signaling pathways. Future Sci OA 10, 2387961 (2024). 10.1080/20565623.2024.2387961

6 Luo, R., Yao, Y., Chen, Z. & Sun, X. An examination of the LPS-TLR4 immune response through the analysis of molecular structures and protein–protein interactions. Cell Communication and Signaling 23, 142 (2025). 10.1186/s12964-025-02149-4

7 O’Neill, L. A. J. & Bowie, A. G. The family of five: TIR-domain-containing adaptors in Toll-like receptor signalling. Nature Reviews Immunology 7, 353–364 (2007). 10.1038/nri2079

8 Kawai, T., Ikegawa, M., Ori, D. & Akira, S. Decoding Toll-like receptors: Recent insights and perspectives in innate immunity. Immunity 57, 649–673 (2024). 10.1016/j.immuni.2024.03.004

9 Fitzgerald, K. A. & Kagan, J. C. Toll-like Receptors and the Control of Immunity. Cell 180, 1044–1066 (2020). 10.1016/j.cell.2020.02.041

10 Wang, K., Huang, H., Zhan, Q., Ding, H. & Li, Y. Toll-like receptors in health and disease. MedComm 5, e549 (2024). 10.1002/mco2.549

11 Hennessy, E. J., Parker, A. E. & O’Neill, L. A. J. Targeting Toll-like receptors: emerging therapeutics? Nature Reviews Drug Discovery 9, 293–307 (2010). 10.1038/nrd3203

12 Thomas, V., Nicholas, J. G., Ashley, M., Bostjan, K. & Stuart, K. Adaptors in Toll-Like Receptor Signaling and their Potential as Therapeutic Targets. Current Drug Targets 13, 1360–1374 (2012). 10.2174/138945012803530260

13 Rajpoot, S. et al. TIRAP in the Mechanism of Inflammation. Frontiers in Immunology Volume 12 - 2021 (2021). 10.3389/fimmu.2021.697588

14 Bernard, N. J. & O’Neill, L. A. Mal, more than a bridge to MyD88. IUBMB Life 65, 777–786 (2013). 10.1002/iub.1201

15 Lin, Z., Lu, J., Zhou, W. & Shen, Y. Structural Insights into TIR Domain Specificity of the Bridging Adaptor Mal in TLR4 Signaling. PLOS ONE 7, e34202 (2012). 10.1371/journal.pone.0034202

16 Bonham, Kevin S. et al. A Promiscuous Lipid-Binding Protein Diversifies the Subcellular Sites of Toll-like Receptor Signal Transduction. Cell 156, 705–716 (2014). 10.1016/j.cell.2014.01.019

17 Lin, S.-C., Lo, Y.-C. & Wu, H. Helical assembly in the MyD88–IRAK4–IRAK2 complex in TLR/IL-1R signalling. Nature 465, 885–890 (2010). 10.1038/nature09121

18 Balka, K. R. & De Nardo, D. Understanding early TLR signaling through the Myddosome. Journal of Leukocyte Biology 105, 339–351 (2018). 10.1002/jlb.Mr0318-096r

19 Kagan, J. C. & Medzhitov, R. Phosphoinositide-Mediated Adaptor Recruitment Controls Toll-like Receptor Signaling. Cell 125, 943–955 (2006). 10.1016/j.cell.2006.03.047

20 Zhao, X. et al. Membrane targeting of TIRAP is negatively regulated by phosphorylation in its phosphoinositide-binding motif. Scientific Reports 7, 43043 (2017). 10.1038/srep43043

21 Patra, M. C. & Choi, S. Insight into Phosphatidylinositol-Dependent Membrane Localization of the Innate Immune Adaptor Protein Toll/Interleukin 1 Receptor Domain-Containing Adaptor Protein. Frontiers in Immunology Volume 9 -2018 (2018). 10.3389/fimmu.2018.00075

22 Bhatt, A. et al. Structural characterization of TIR-domain signalosomes through a combination of structural biology approaches. IUCrJ 11, 695–707 (2024). doi:10.1107/S2052252524007693

23 Patra, M. C. & Choi, S. Insight into Phosphatidylinositol-Dependent Membrane Localization of the Innate Immune Adaptor Protein Toll/Interleukin 1 Receptor Domain-Containing Adaptor Protein. Front Immunol 9, 75 (2018). 10.3389/fimmu.2018.00075

24 Ve, T. et al. Structural basis of TIR-domain-assembly formation in MAL- and MyD88-dependent TLR4 signaling. Nature Structural & Molecular Biology 24, 743–751 (2017). 10.1038/nsmb.3444

25 Clabbers, M. T. B. et al. MyD88 TIR domain higher-order assembly interactions revealed by microcrystal electron diffraction and serial femtosecond crystallography. Nature Communications 12, 2578 (2021). 10.1038/s41467-021-22590-6

26 Manik, M. K. et al. Structural basis for TIR domain–mediated innate immune signaling by Toll-like receptor adaptors TRIF and TRAM. Proceedings of the National Academy of Sciences 122, e2418988122 (2025). doi:10.1073/pnas.2418988122

27 Nanson, J. D., Kobe, B. & Ve, T. Death, TIR, and RHIM: Self-assembling domains involved in innate immunity and cell-death signaling. J Leukoc Biol 105, 363–375 (2019). 10.1002/jlb.Mr0318-123r

28 Kobe, B. et al. Signalling by co-operative higher-order assembly formation: linking evidence at molecular and cellular levels. Biochemical Journal 482, 275–294 (2025). 10.1042/bcj20220094

29 Nimma, S. et al. Structural Evolution of TIR-Domain Signalosomes. Frontiers in Immunology Volume 12 - 2021 (2021). 10.3389/fimmu.2021.784484

30 Stark, H. GraFix: stabilization of fragile macromolecular complexes for single particle cryo-EM. Methods Enzymol 481, 109–126 (2010). 10.1016/s0076-6879(10)81005-5

31 Lata, S., Gavutis, M., Tampé, R. & Piehler, J. Specific and Stable Fluorescence Labeling of Histidine-Tagged Proteins for Dissecting Multi-Protein Complex Formation. Journal of the American Chemical Society 128, 2365–2372 (2006). 10.1021/ja0563105

32 Bartoschik, T. et al. Near-native, site-specific and purification-free protein labeling for quantitative protein interaction analysis by MicroScale Thermophoresis. Scientific Reports 8, 4977 (2018). 10.1038/s41598-018-23154-3

33 Vajjhala, P. R., Ve, T., Bentham, A., Stacey, K. J. & Kobe, B. The molecular mechanisms of signaling by cooperative assembly formation in innate immunity pathways. Molecular Immunology 86, 23–37 (2017). 10.1016/j.molimm.2017.02.012

34 Schnitzbauer, J., Strauss, M. T., Schlichthaerle, T., Schueder, F. & Jungmann, R. Super-resolution microscopy with DNA-PAINT. Nature Protocols 12, 1198–1228 (2017). 10.1038/nprot.2017.024

35 Nieves, D. J., Gaus, K. & Baker, M. A. B. DNA-Based Super-Resolution Microscopy: DNA-PAINT. Genes 9, 621 (2018).

36 Kappelhoff, S. et al. Structure and regulation of GSDMD pores at the plasma membrane of pyroptotic cells. bioRxiv, 2023.2010.2024.563742 (2023). 10.1101/2023.10.24.563742

37 Lelek, M. et al. Single-molecule localization microscopy. Nat Rev Methods Primers 1 (2021). 10.1038/s43586-021-00038-x

38 Früh, S. M. et al. Site-Specifically-Labeled Antibodies for Super-Resolution Microscopy Reveal In Situ Linkage Errors. ACS Nano 15, 12161–12170 (2021). 10.1021/acsnano.1c03677

39 Li, M. & Yu, Y. Innate immune receptor clustering and its role in immune regulation. J Cell Sci 134 (2021). 10.1242/jcs.249318

40 Heuer, L. S. et al. Higher order receptor clustering due to the IgG3 subclass is necessary for TLR4 signaling and tolerance induction by novel human anti-TLR4 antibodies. MAbs 17, 2515415 (2025). 10.1080/19420862.2025.2515415

41 Felker, A. et al. A Versatile Toolbox for Nanoscale Interrogation of Multiprotein Assemblies inside Living Cells. bioRxiv, 2025.2004.2030.651189 (2025). 10.1101/2025.04.30.651189

42 Philippi, M. et al. Biofunctional Nanodot Arrays in Living Cells Uncover Synergistic Co-Condensation of Wnt Signalodroplets. Small 18, 2203723 (2022). 10.1002/smll.202203723

43 Schmidt, M. et al. Capillary Nanostamping with Spongy Mesoporous Silica Stamps. Advanced Functional Materials 28, 1800700 (2018). 10.1002/adfm.201800700

44 Brauchle, M. et al. Protein interference applications in cellular and developmental biology using DARPins that recognize GFP and mCherry. Biol Open 3, 1252–1261 (2014). 10.1242/bio.201410041

45 Del Val, E., Fernández-Vega, A., Molina, M. & Cid, V. J. MyD88 polymerization and association to cellular membranes in a yeast heterologous model. Cell Mol Life Sci 82, 288 (2025). 10.1007/s00018-025-05827-1

46 Kasai, K. et al. From Monomers to Oligomers: Structural Mechanism of Receptor-Triggered MyD88 Assembly in Innate Immune Signaling. bioRxiv, 2024.2009.2013.612588 (2024). 10.1101/2024.09.13.612588

47 Cao, F., Deliz-Aguirre, R., Gerpott, F. H., Ziska, E. & Taylor, M. J. Myddosome clustering in IL-1 receptor signaling regulates the formation of an NF-kB activating signalosome. EMBO Rep 24, e57233 (2023). 10.15252/embr.202357233

48 Tompa, P., Schad, E., Tantos, A. & Kalmar, L. Intrinsically disordered proteins: emerging interaction specialists. Current Opinion in Structural Biology 35, 49–59 (2015). 10.1016/j.sbi.2015.08.009

49 Wu, H. & Fuxreiter, M. The Structure and Dynamics of Higher-Order Assemblies: Amyloids, Signalosomes, and Granules. Cell 165, 1055–1066 (2016). 10.1016/j.cell.2016.05.004

50 Latty, S. L. et al. Activation of Toll-like receptors nucleates assembly of the MyDDosome signaling hub. eLife 7, e31377 (2018). 10.7554/eLife.31377

51 Moncrieffe, M. C. et al. MyD88 Death-Domain Oligomerization Determines Myddosome Architecture: Implications for Toll-like Receptor Signaling. Structure 28, 281–289.e283 (2020). 10.1016/j.str.2020.01.003

52 Januliene, D. & Moeller, A. Single-Particle Cryo-EM of Membrane Proteins. Methods Mol Biol 2302, 153–178 (2021). 10.1007/978-1-0716-1394-8_9

53 Punjani, A., Rubinstein, J. L., Fleet, D. J. & Brubaker, M. A. cryoSPARC: algorithms for rapid unsupervised cryo-EM structure determination. Nat Methods 14, 290–296 (2017). 10.1038/nmeth.4169

54 Liebschner, D. et al. Macromolecular structure determination using X-rays, neutrons and electrons: recent developments in Phenix. Acta Crystallogr D Struct Biol 75, 861–877 (2019). 10.1107/s2059798319011471

55 Pettersen, E. F. et al. UCSF ChimeraX: Structure visualization for researchers, educators, and developers. Protein Sci 30, 70–82 (2021). 10.1002/pro.3943

56 Emsley, P. & Cowtan, K. Coot: model-building tools for molecular graphics. Acta Crystallogr D Biol Crystallogr 60, 2126–2132 (2004). 10.1107/s0907444904019158

57 Chen, V. B. et al. MolProbity: all-atom structure validation for macromolecular crystallography. Acta Crystallogr D Biol Crystallogr 66, 12–21 (2010). 10.1107/s0907444909042073

58 Han, X., Yang, Y., Zhao, F., Zhang, T. & Yu, X. An improved protein lipid overlay assay for studying lipid–protein interactions. Plant Methods 16, 33 (2020). 10.1186/s13007-020-00578-5

59 Schindelin, J., et al. Fiji: an open-source platform for biological-image analysis. Nat Methods 9, 676–682 (2012). 10.1038/nmeth.2019

60 Bentea, L., Watzky, M. A. & Finke, R. G. Sigmoidal Nucleation and Growth Curves Across Nature Fit by the Finke–Watzky Model of Slow Continuous Nucleation and Autocatalytic Growth: Explicit Formulas for the Lag and Growth Times Plus Other Key Insights. The Journal of Physical Chemistry C 121, 5302–5312 (2017). 10.1021/acs.jpcc.6b12021

61 Arganda-Carreras, I. et al. Trainable Weka Segmentation: a machine learning tool for microscopy pixel classification. Bioinformatics 33, 2424–2426 (2017). 10.1093/bioinformatics/btx180

62 Wedeking, T. et al. Spatiotemporally Controlled Reorganization of Signaling Complexes in the Plasma Membrane of Living Cells. Small 11, 5912–5918 (2015). 10.1002/smll.201502132

63 Mortensen, K. I., Churchman, L. S., Spudich, J. A. & Flyvbjerg, H. Optimized localization analysis for single-molecule tracking and super-resolution microscopy. Nature Methods 7, 377–381 (2010). 10.1038/nmeth.1447

64 Wang, Y. et al. Localization events-based sample drift correction for localization microscopy with redundant cross-correlation algorithm. Opt. Express 22, 15982–15991 (2014). 10.1364/OE.22.015982

65 Margheritis, E. et al. Gasdermin D cysteine residues synergistically control its palmitoylation-mediated membrane targeting and assembly. The EMBO Journal 43, 4274–4297 (2024). 10.1038/s44318-024-00190-6

66 Philippi, M. et al. Close-packed silane nanodot arrays by capillary nanostamping coupled with heterocyclic silane ring opening. RSC Advances 9, 24742–24750 (2019). 10.1039/C9RA03440D

67 Krissinel, E. & Henrick, K. Inference of macromolecular assemblies from crystalline state. J Mol Biol 372, 774–797 (2007). 10.1016/j.jmb.2007.05.022

68 Laskowski, R. A. & Swindells, M. B. LigPlot+: multiple ligand-protein interaction diagrams for drug discovery. J Chem Inf Model 51, 2778–2786 (2011). 10.1021/ci200227u

