## Supplementary Figures and Tables for "Ultrastructural organization and dynamics of TIRAP filaments"

#### Affiliations

#### This PDF file includes:

Supplementary Figures 1 to 10

Supplementary Tables 1 to 3

Descriptions of Movie Files S1 to S6

#### Supplementary Figures

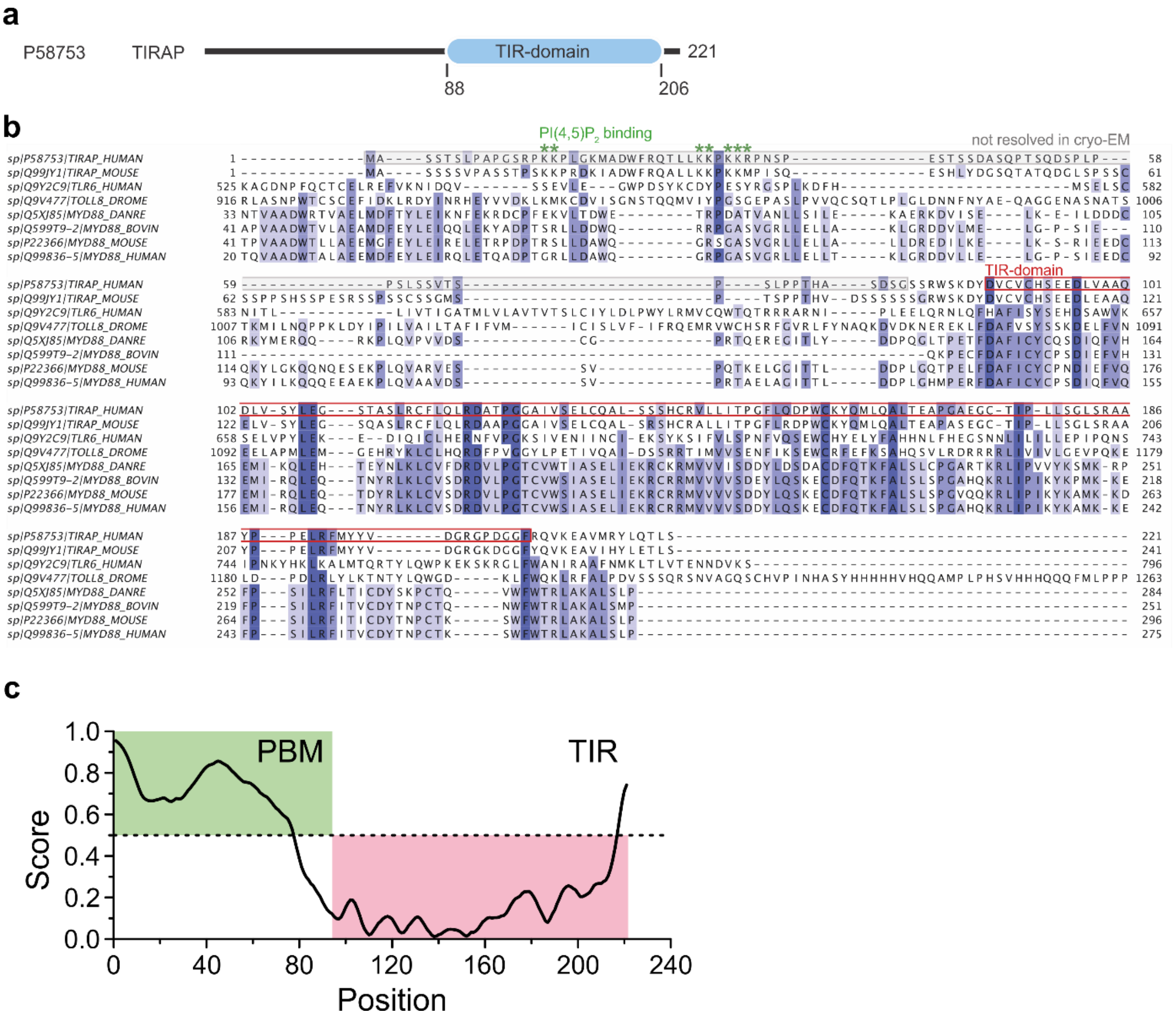

**Supplementary Fig. 1: Domain architecture and sequence analysis of TIRAP.** **a** Domain organization of human TIRAP (P58753) with PBM (residues 1-88) and TIR domain (residues 88-221) indicated. **b** Multiple sequence alignment (MSA) of TIRAP orthologs showing conservation of the PBM region (green box highlighting PI(4,5)P$_2$ binding residues) and TIR domain (red box). Sequences are color coded by conservation with a cut-off at 30 %. Sequences were selected by highest sequence identity hits from protein BLAST of selected model organisms. MSAs were prepared with Jalview[1]. Key di-lysine clusters (K15-K16, K31-K32, K35-K36) are marked with green asterisks[2]. Note: TIR domain boundaries are shaded in red; N-terminal region remains largely unresolved in cryo-EM (marked "not resolved in cryo-EM"). **c** Disorder prediction for human TIRAP using IUPred3[3]. The PBM exhibits high disorder scores (> 0.5, green shading), while the TIR domain shows low disorder (pink shading). Dashed line indicates disorder threshold (0.5).

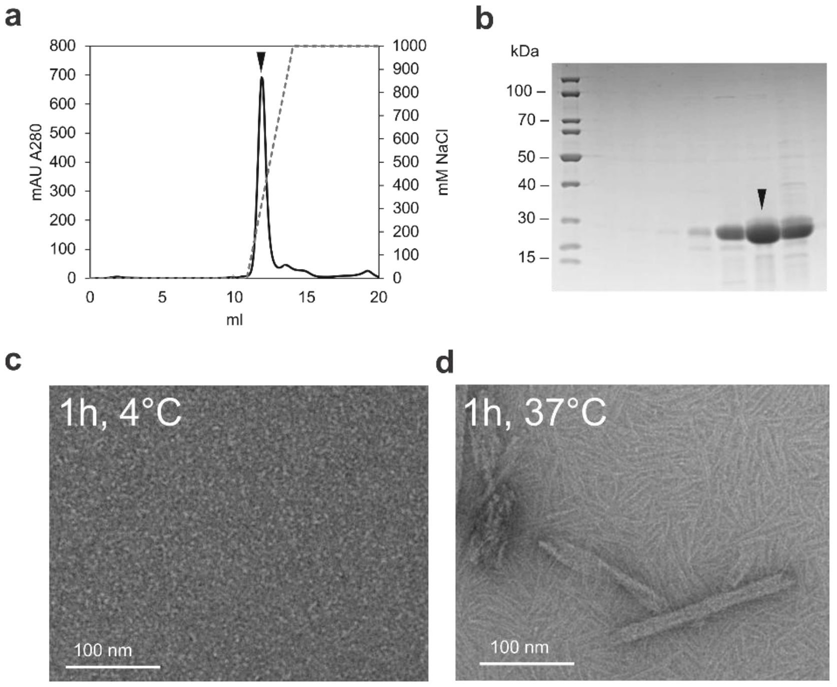

**Supplementary Fig. 2: Protein purification, and temperature-dependent assembly.** **a** Ion-exchange chromatography profile of TIRAP. Black arrow corresponds to indicated SDS-PAGE fraction. **b** Coomassie-blue stained SDS-PAGE gel of purified TIRAP. Black arrow corresponds to IEX fraction. **c-d** Representative micrographs from negative-stain TEM. Control after 1 h incubation at 4°C (c) and 37°C (d). Scale bar: 100 nm.

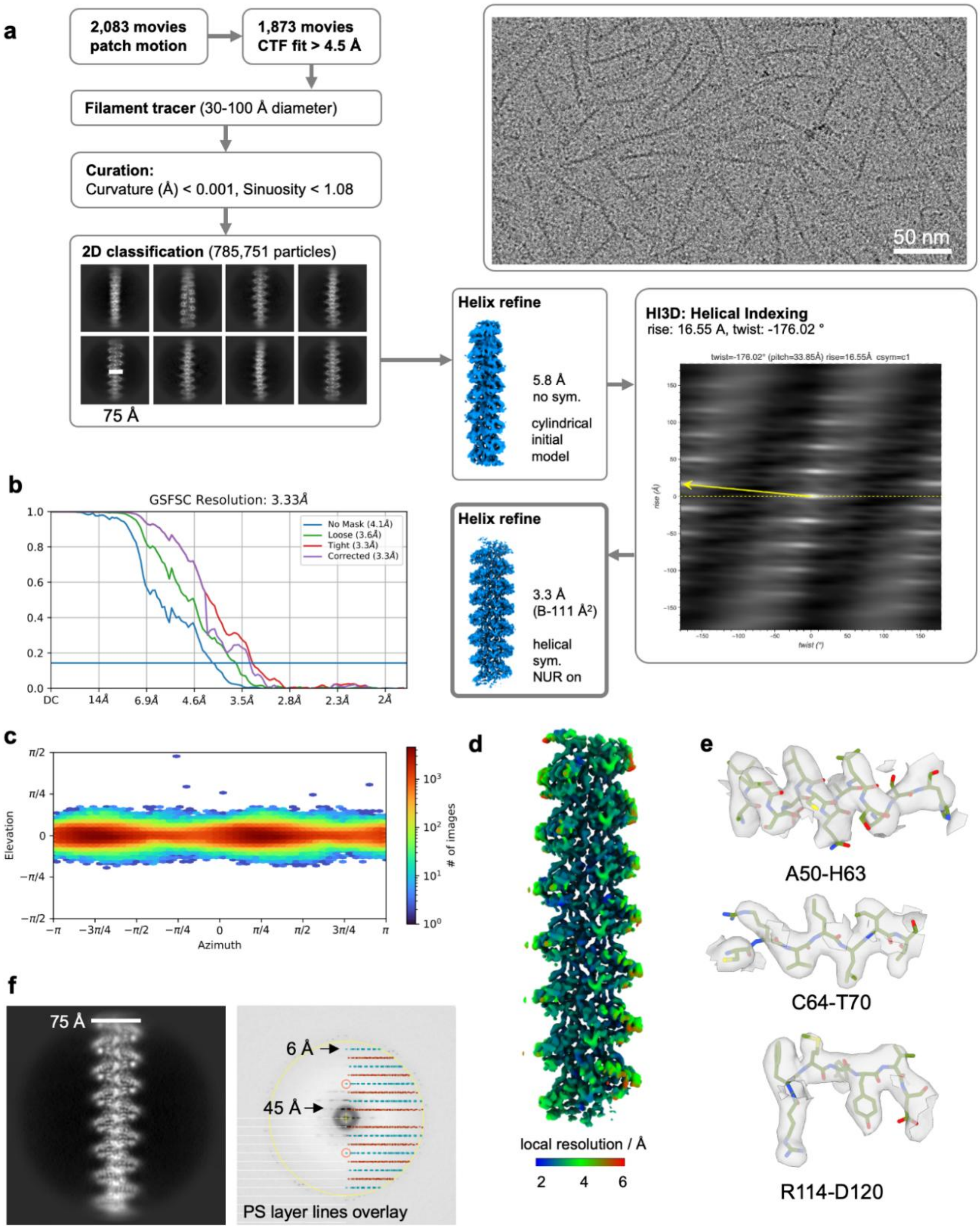

**Supplementary Fig. 3: Cryo-EM data processing and helical reconstruction.** **a** Processing workflow for the TIRAP filament in its apo state. All processing steps were performed in cryoSPARC. Representative cryo-EM micrograph and 2D-class averages are given. 50 nm scale bar in micrograph and 70 Å scale bar in the 2D classes. **b** Gold-standard Fourier shell correlation (FSC) curves showing global resolution of 3.3 Å at FSC = 0.143. Curves shown: no mask (blue, 4.1 Å), loose mask (green, 3.6 Å), tight mask (red, 3.3 Å), and phase-randomization corrected (purple, 3.3 Å). **c** Angular particle distribution plot showing uniform angular coverage in helical reconstruction. **d** Local resolution map of the TIRAP helical filament color-coded from 2 Å (blue) to 6 Å (red). **e** Representative cryo-EM density for three regions of the map showing side-chain density: A50-H63, C64-T70, and R114-D120. **f** 2D class average and power spectrum (PS). The overlayed layer lines were annotated using Helixplorer[4].

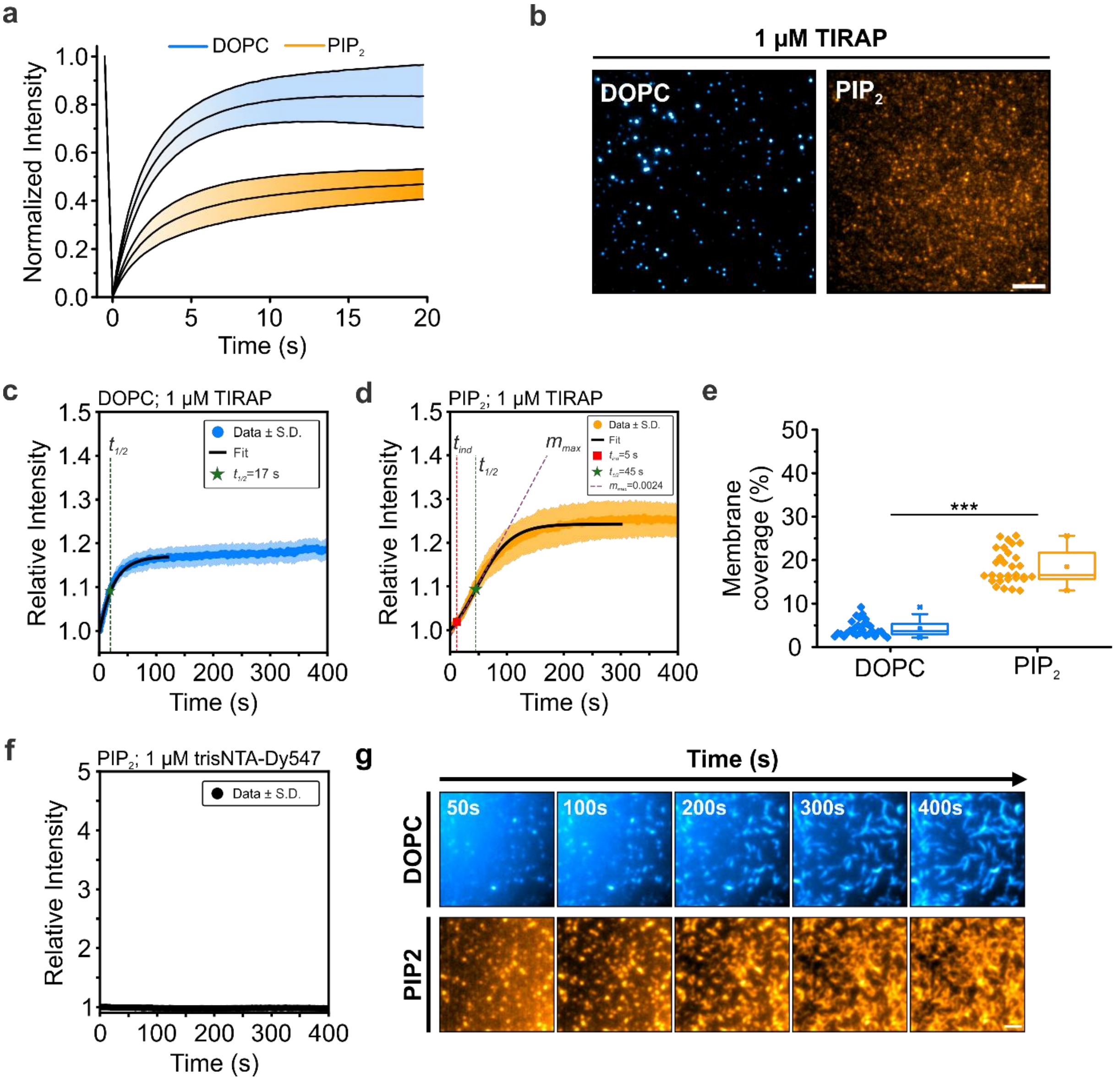

**Supplementary Fig. 4: Concentration- and lipid-dependent kinetics of TIRAP membrane recruitment and assembly. a** Fluorescence recovery after photobleaching (FRAP) recovery curves from DOPC (100 mol%, blue) and DOPC:$PIP_2$ (95:5 mol%, orange) containing SLBs. **b** Representative TIRF images (1 µM TIRAP) on DOPC (blue) and $PIP_2$-containing bilayers (orange). Scale bar: 5 µm. **c** Kinetics of 1 µM TIRAP on DOPC bilayers. Blue: data ± s.d.; black line: exponential fit. Key parameters are indicated. **d** Kinetics of 1 µM TIRAP on $PIP_2$ bilayers. Orange: data ± s.d.; black line: Finke-Watzky[5] fit. Key parameters are indicated. **e** Membrane coverage (%) at 1 µM TIRAP. Box plot indicates data distribution of the second and third quartiles (box), median (line), mean (square), and 1.5x interquartile range (whiskers). Each dot represent individual regions of interest (n > 25 ROIs per condition, 2 independent experiments). ***$p < 0.001$, two-sample Kolmogorov-Smirnov test. **f** Nonspecific membrane binding of trisNTA-Dy547 (1 µM) without protein incubation. Black: data ± s.d. **g** Representative time-lapse TIRF images (10 µM TIRAP) on DOPC (blue, top) and $PIP_2$-containing bilayers (orange, bottom) showing raw fluorescence images. Scale bar: 2 µm.

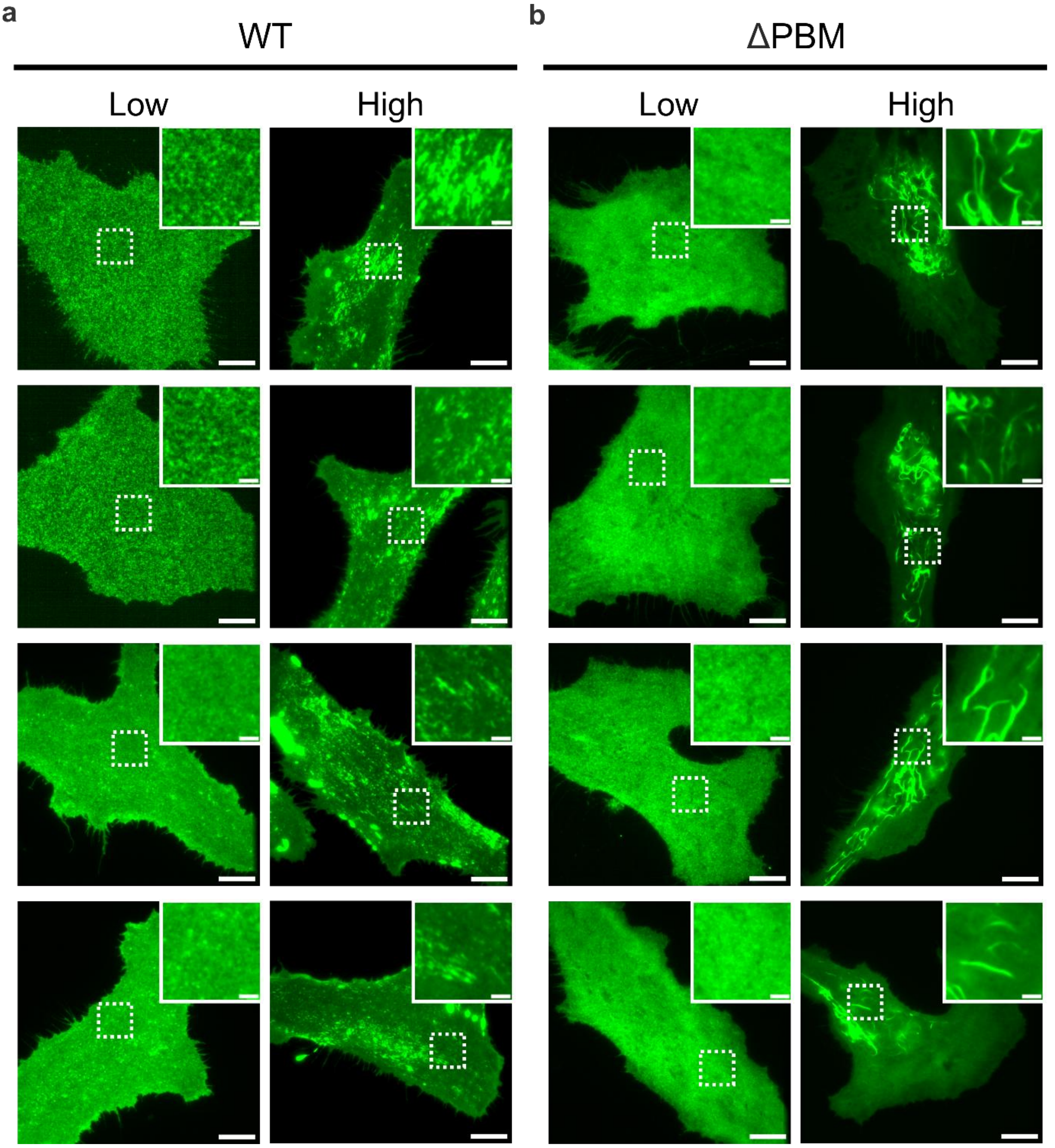

**Supplementary Fig. 5: Plasma membrane localization of TIRAP variants at different expression levels. a, b** Representative TIRF images of HeLa cells expressing TIRAP-wt (**a**) or TIRAP-ΔPBM (**b**) at low (left) and high (right) PM intensity. Multiple cells shown per condition (rows). Insets show magnified regions indicated by white dashed boxes. Scale bars: 10 μm; insets, 2 μm.

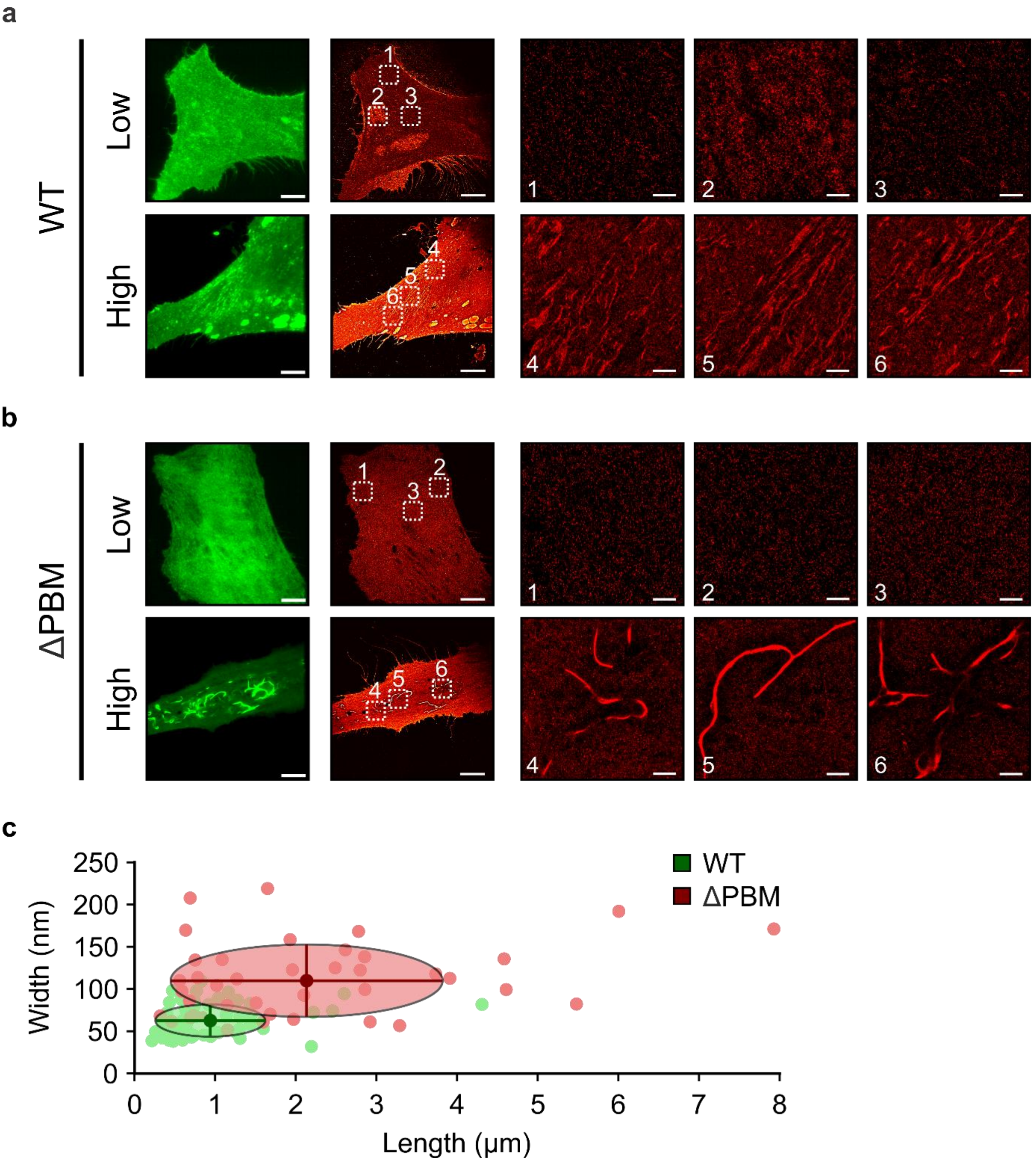

**Supplementary Fig. 6: DNA-PAINT images and analysis of TIRAP filament morphology. a, b** Representative TIRF (left, green) and DNA-PAINT (right, red) images for TIRAP-wt (**a**) and TIRAP-ΔPBM (**b**) at low (top) and high (bottom) PM intensity. White dashed boxes indicate regions of interest shown as magnified insets (ROIs 1-6). Scale bars: overview images: 10 µm; DNA-PAINT insets: 1 µm. **c** Scatter plot of filament width versus length for TIRAP-wt (green) and TIRAP-ΔPBM (red) at high PM intensity. Ellipses represent 95% confidence intervals.

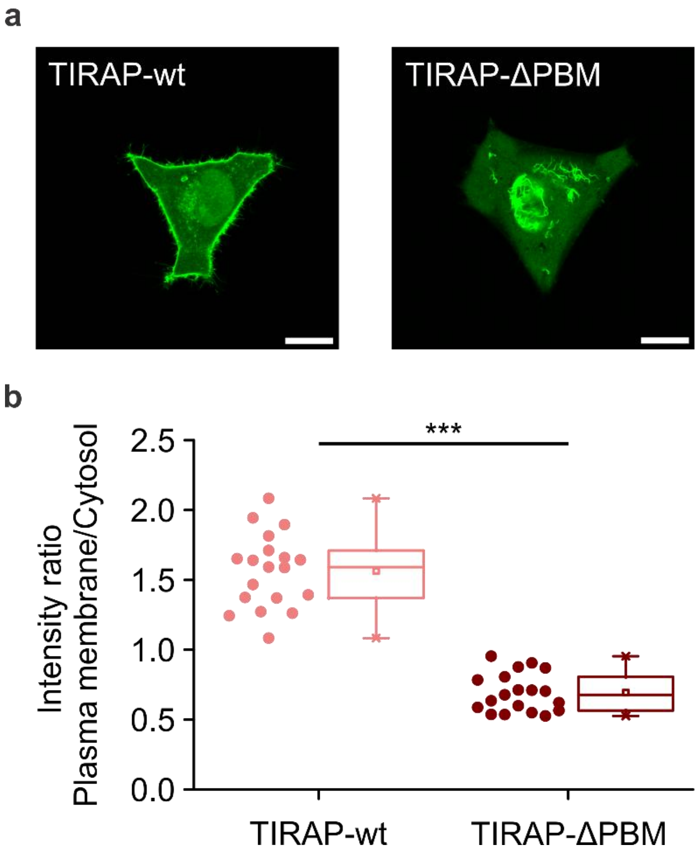

**Supplementary Fig. 7: Plasma membrane localization of TIRAP variants.** **a** Representative confocal microscopy images of HeLa cells expressing mEGFP-TIRAP-wt (left) or mEGFP-TIRAP-ΔPBM (right). Scale bars: 10 µm. **b** Quantification of plasma membrane-to-cytosol fluorescence intensity ratios (TIRAP-wt: n = 19 cells; TIRAP-ΔPBM: n = 19 cells; ***$p < 0.001$, two-sample Kolmogorov-Smirnov test). Box plot indicates data distribution of the second and third quartiles (box), median (line), mean (square), and 1.5x interquartile range (whiskers).

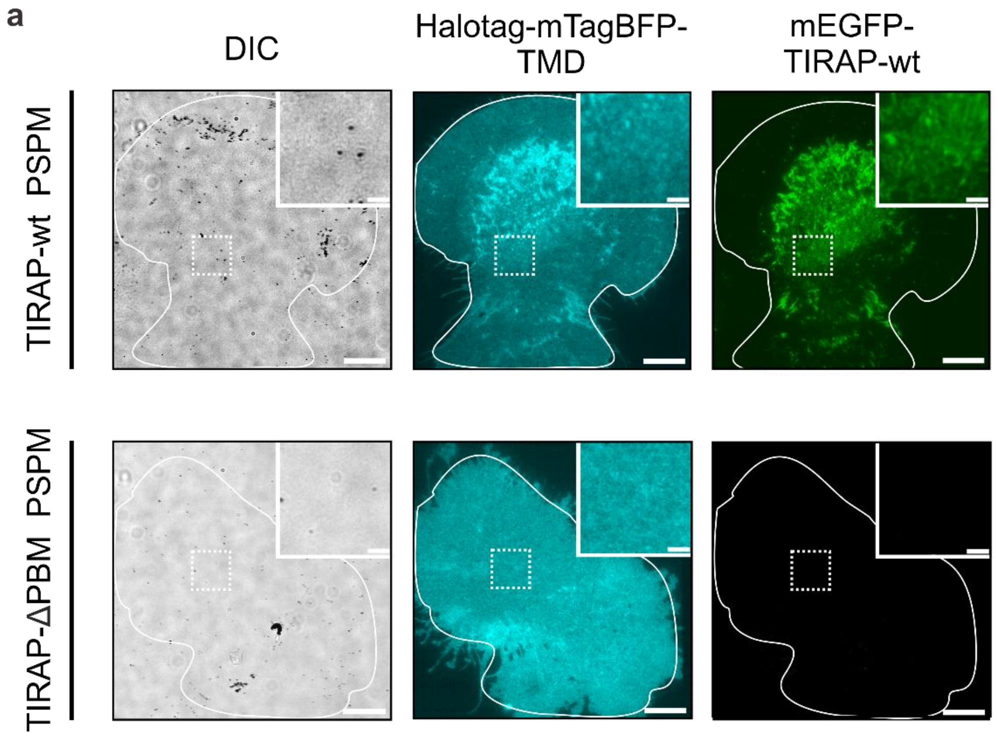

**Supplementary Fig. 8: Validation of PSPM preparation for TIRAP variants. a** Representative TIRF images illustrating successful PSPM generation. Top row: PSPM from TIRAP-wt-expressing cell. Bottom row: PSPM from TIRAP-ΔPBM-expressing cell. Left: DIC image confirming absence of intact cell body. Middle: HaloTag-mTagBFP-TMD showing homogeneous membrane distribution. Right: mEGFP-TIRAP fluorescence signal. White outlines indicate membrane boundaries; dashed boxes indicate magnified regions shown as insets. Images were contrasted with same parameters. Scale bars: overview, 10 μm; insets, 2 μm.

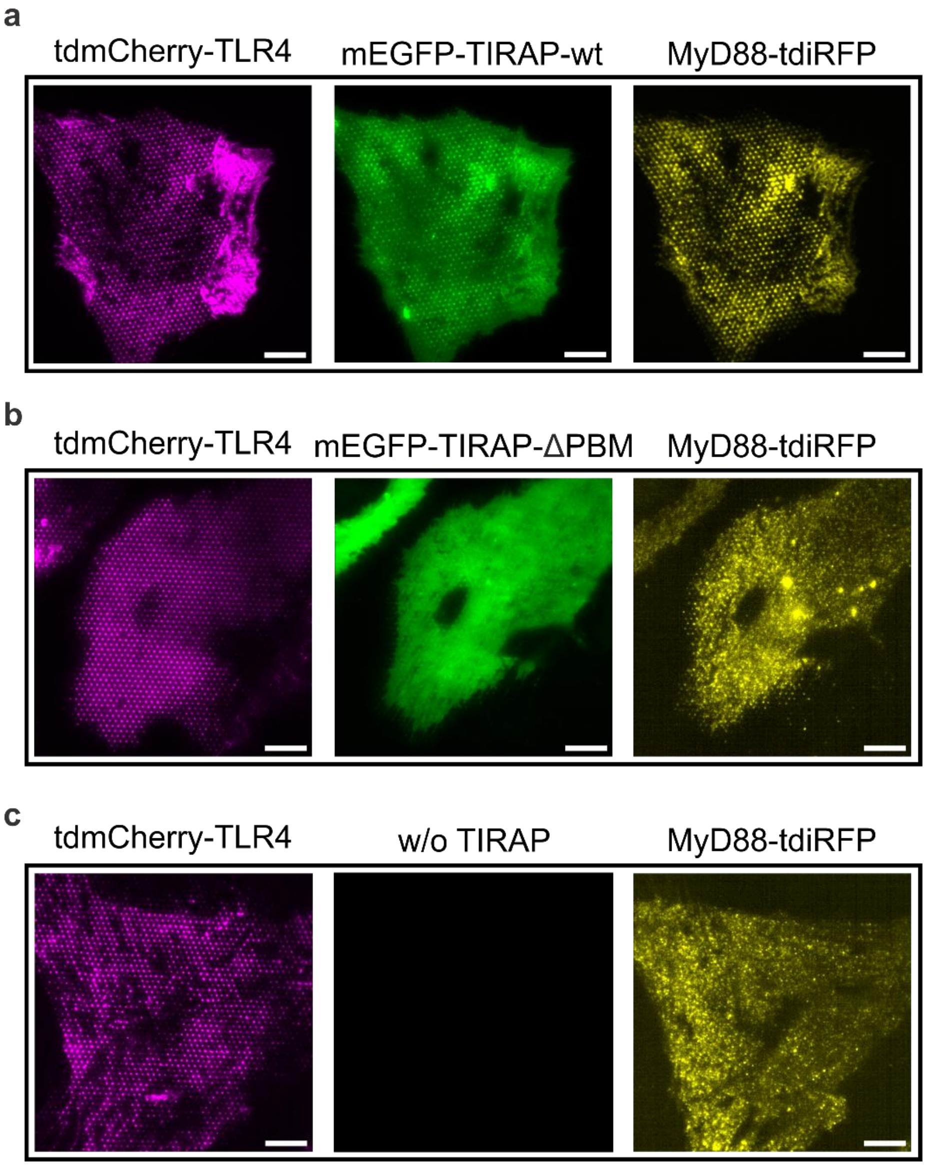

**Supplementary Fig. 9: Representative TIRF microscopy images illustrating recruitment of TIRAP and MyD88 to immobilized TLR4 receptors. a-c** Representative TIRF microscopy images showing tdmCherry-TLR4 (magenta), mEGFP-TIRAP (green), and MyD88-tdiRFP (yellow) for TIRAP-wt (**a**), TIRAP-ΔPBM (**b**), and cells without TIRAP (**c**). Scale bar: 5 µm.

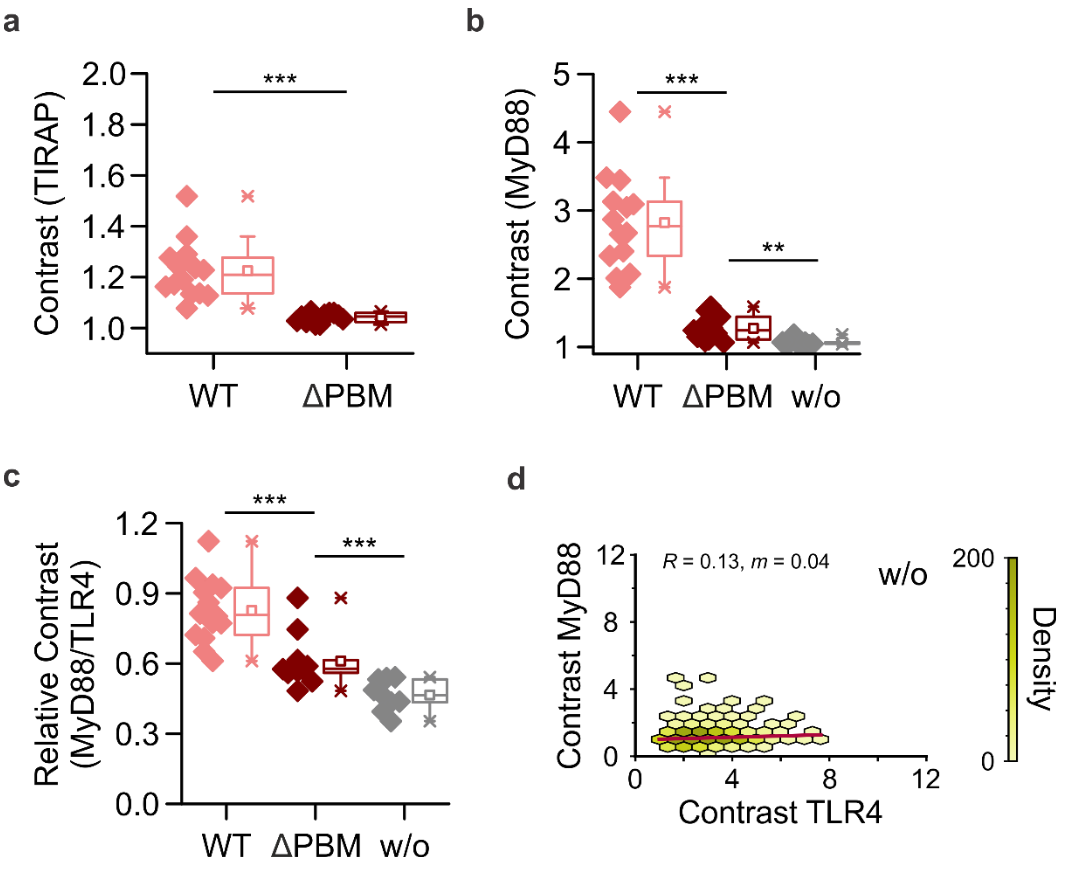

**Supplementary Fig. 10: Quantitative analysis of adaptor recruitment. a** Mean TIRAP contrast values per cell for TIRAP-wt (n = 14 cells) and TIRAP-ΔPBM (n = 11 cells). **b** Mean MyD88 contrast values per cell for TIRAP-wt (n = 14 cells), TIRAP-ΔPBM (n = 11 cells), and w/o TIRAP (n = 9 cells). **c** Mean relative contrast (MyD88/TLR4) values per cell for TIRAP-wt (n = 14 cells), TIRAP-ΔPBM (n = 11 cells), and w/o TIRAP (n = 9 cells). Box plots indicate data distribution of the second and third quartiles (box), median (line), mean (square), and 1.5x interquartile range (whiskers). $^{***}p < 0.001$, $^{**}p < 0.01$ (Two-sample Kolmogorov-Smirnov test). **d** Hexbin density scatter plot showing correlation between TLR4 and MyD88 contrast for cells without TIRAP expression (n > 4.000 nanodots from 9 cells). Pearson correlation coefficient (*R*) and linear regression slope (*m*) are indicated.

### Supplementary Tables

#### Supplementary Table 1: Cryo-EM data collection, refinement and validation statistics.

| | ***Hs*TIRAP (PDB 9FQM, EMD-50657)** |
|---|---|
| **Data collection and processing** | |
| **Microscope** | TFS Glacios |
| **Voltage (keV)** | 200 |
| **Camera** | TFS Falcon 4 |
| **Energy filter** | TFS Selectris |
| **Slit width eV** | 10 |
| **Magnification (nominal)** | 130,000 |
| **Pixel size (Å/px)** | 0.945 |
| **Defocus range (µm)** | 0.8-2.0 |
| **Total exposure ($e^-$/Å$^2$)** | 50 |
| **Automation software** | *EPU 2.9* |
| **Processing software** | *CryoSPARC* 4 |
| **Micrographs collected** | 2,083 |
| **Micrographs used** | 1,172 |
| **Final particle images** | 785,751 |
| **Helical symmetry parameters**<br>**Twist**<br>**Rise** | <br>-176°<br>16.6 Å |
| **Resolution (global) (Å) FSC 0.134** | 3.3 |
| **Map-sharpening *B* factor (Å$^2$)** | – 111 |
| **Map-sharpening method** | Global *B* factor |
| **Refinement package** | *phenix.real_space_refine* |
| **Model Composition** | |
| **Chains** | 8 |
| **Non-H atoms** | 8,808 |
| **Protein Residues** | 1,144 |
| **Model refinement** | |
| **Model-map scores** | |
| **FSC model (0.5)** | 4.0 |
| **Av. grouped B factors (Å$^2$)** | |
| **Protein residues** | 83.94 |
| **R.m.s.d. from ideal values** | |
| **Bond lenghths (Å)** | 0.003 |
| **Bond angles (°)** | 0.561 |
| **Validation** | |
| ***MolProbity* score** | 1.71 |
| **CaBLAM outliers (%)** | 1.17 |
| **Clashscore** | 10.48 |
| **Poor rotamers (%)** | 0.00 |
| **$C^\beta$ outliers (%)** | 0.00 |
| **Ramachandran plot** | |
| **Favored (%)** | 96.99 |
| **Allowed (%)** | 3.01 |
| **Outliers (%)** | 0.00 |

#### Supplementary Table 2: Kinetic parameters for TIRAP assembly on supported lipid bilayers.

Concentration-dependent assembly kinetics determined by exponential (DOPC, bottom) or Finke-Watzky ($PIP_2$, top) fitting.

| c (µM) | $t_{ind}$ (s) | $t_{1/2}$ (s) | $m_{max}$ | $k_1$ ($s^{-1}$) | $k_2$ ($M^{-1}s^{-1}$) | Fit range (s) | $R^2$ | $I_{rel,max}$ |
|---|---|---|---|---|---|---|---|---|
| **1** | 4.71 | 44.98 | $2.44\times10^{-3}$ | $(6.11\pm0.39)\times10^{-3}$ | $(1.10\pm0.12)\times10^{-1}$ | 1-300 | 0.989 | 1.25 |
| **10** | 68.37 | 129.94 | $1.84\times10^{-2}$ | $(1.25\pm0.02)\times10^{-3}$ | $(6.21\pm0.12)\times10^{-3}$ | 1-310 | 0.999 | 4.21 |

| c (µM) | $t_{1/2}$ (s) | k ($s^{-1}$) | Fit range (s) | $R^2$ | $I_{rel,max}$ |
|---|---|---|---|---|---|
| **1** | 17.19 | $(4.03\pm0.18)\times10^{-2}$ | 1-120 | 0.992 | 1.17 |
| **10** | 14.97 | $(4.63\pm0.16)\times10^{-2}$ | 1-120 | 0.982 | 1.99 |

c: TIRAP concentration; $t_{ind}$: induction time; $t_{1/2}$: half-time to 50% plateau; $m_{max}$: maximum assembly slope at half-time; $k_1$: nucleation rate constant; $k_2$: growth rate constant; $k$: exponential rate constant; $I_{rel,max}$ = relative plateau intensity normalized to initial fluorescence; Values represent mean ± s.e.m. from n = 12 independent regions per condition across 2 independent experiments.

#### Supplementary Table 3: Recombinant DNA.

| Denomination | Plasmid name | Source |
|---|---|---|
| **$His_6$-TIRAP** | pet28a-6xHis-Myc-TEV-hTIRAP | This study |
| **aALFAnb-td2m22** | pet21a-antimCherryDARPin2m22-linker-antiALFA nanobody-linker-antimCherryDARPin2m22-6xHis | Felker et al.[6] |
| **TIRAP-wt** | pSems-mEGFP-hTIRAP | This study |
| **TIRAP-ΔPBM** | pSems-mEGFP-hTIRAP(S80-221) | This study |
| **HaloTag-mTagBFP-TMD** | pDisplay-HaloTag-mTagBFP-TMD-GSlinker | Kappelhoff et al.[7] |
| **tdmCherry-TLR4** | pSems-leader-2xmCherry-hTLR4 | This study |
| **MyD88-tdiRFP** | pSems-mMyD88-tdiRFP | This study |

Linker sequence:SSGGS

TMD sequence: ASALAALAALAALAALAALAALAKSSRL

#### References

1 Waterhouse, A. M., Procter, J. B., Martin, D. M., Clamp, M. & Barton, G. J. Jalview Version 2--a multiple sequence alignment editor and analysis workbench. *Bioinformatics* **25**, 1189-1191 (2009). https://doi.org:10.1093/bioinformatics/btp033

2 Kagan, J. C. & Medzhitov, R. Phosphoinositide-Mediated Adaptor Recruitment Controls Toll-like Receptor Signaling. *Cell* **125**, 943-955 (2006). https://doi.org:https://doi.org/10.1016/j.cell.2006.03.047

3 Erdős, G., Pajkos, M. & Dosztányi, Z. IUPred3: prediction of protein disorder enhanced with unambiguous experimental annotation and visualization of evolutionary conservation. *Nucleic Acids Research* **49**, W297-W303 (2021). https://doi.org:10.1093/nar/gkab408

4 Egelman, E. H. Helical reconstruction, again. *Current Opinion in Structural Biology* **85**, 102788 (2024). https://doi.org:https://doi.org/10.1016/j.sbi.2024.102788

5 Bentea, L., Watzky, M. A. & Finke, R. G. Sigmoidal Nucleation and Growth Curves Across Nature Fit by the Finke–Watzky Model of Slow Continuous Nucleation and Autocatalytic Growth: Explicit Formulas for the Lag and Growth Times Plus Other Key Insights. *The Journal of Physical Chemistry C* **121**, 5302-5312 (2017). https://doi.org:10.1021/acs.jpcc.6b12021

6 Felker, A. *et al.* A Versatile Toolbox for Nanoscale Interrogation of Multiprotein Assemblies inside Living Cells. *bioRxiv*, 2025.2004.2030.651189 (2025). https://doi.org:10.1101/2025.04.30.651189

7 Kappelhoff, S. *et al.* Structure and regulation of GSDMD pores at the plasma membrane of pyroptotic cells. *bioRxiv*, 2023.2010.2024.563742 (2023). https://doi.org:10.1101/2023.10.24.563742

#### Supplementary Movies

**File Name:** Supplementary Movie S1

**Description: Time-lapse TIRF microscopy of TIRAP (1 µM) binding on DOPC membranes.** $His_6$-TIRAP (1 µM) labeled with 100 nM Dy547-trisNTA was applied to supported lipid bilayers composed of 100 mol% DOPC and imaged at 37°C. Movie shows 50-frame temporal binning of fluorescence signal. Scale bar: 2 µm.

**File Name:** Supplementary Movie S2

**Description: Time-lapse TIRF microscopy of TIRAP (10 µM) binding on DOPC membranes.** $His_6$-TIRAP (10 µM) labeled with 1 µM Dy547-trisNTA was applied to supported lipid bilayers composed of 100 mol% DOPC and imaged at 37°C. Movie shows 50-frame temporal binning of fluorescence signal. Scale bar: 2 µm.

**File Name:** Supplementary Movie S3

**Description: Skeletonized representation of time-lapse TIRF microscopy of TIRAP (10 µM) binding on DOPC membranes.** Postprocessed time-lapse data from Supplementary Movie S2 showing segmented and skeletonized filament structures of 10 µM TIRAP on 100 mol% DOPC bilayers. Movie shows 50-frame temporal binning. Scale bar: 2 µm.

**File Name:** Supplementary Movie S4

**Description: Time-lapse TIRF microscopy of TIRAP (1 µM) binding on DOPC:$PIP_2$ membranes.** $His_6$-TIRAP (1 µM) labeled with 100 nM Dy547-trisNTA was applied to supported lipid bilayers composed of DOPC:$PIP_2$ (95:5 mol%) and imaged at 37°C. Movie shows 50-frame temporal binning of fluorescence signal. Scale bar: 2 µm.

**File Name:** Supplementary Movie S5

**Description: Time-lapse TIRF microscopy of TIRAP (10 µM) binding on DOPC:$PIP_2$ membranes.** $His_6$-TIRAP (10 µM) labeled with 1 µM Dy547-trisNTA was applied to supported lipid bilayers composed of DOPC:$PIP_2$ (95:5 mol%) and imaged at 37°C. Movie shows 50-frame temporal binning of fluorescence signal. Scale bar: 2 µm.

**File Name:** Supplementary Movie S6

**Description: Skeletonized representation of time-lapse TIRF microscopy of TIRAP (10 µM) binding on DOPC:PIP$_2$ membranes.** Postprocessed time-lapse data from Supplementary Movie S5 showing segmented and skeletonized filament structures of 10 µM TIRAP on DOPC:PIP$_2$ (95:5 mol%) bilayers. Movie shows 50-frame temporal binning. Scale bar: 2 µm.
